# A projection atlas of excitatory and inhibitory inputs to the preBötzinger Complex: substrates for multimodal breathing control

**DOI:** 10.64898/2026.02.02.703188

**Authors:** Elora K Reilly, Aleeza Kadri, Joseph W Arthurs, Jonathan R Sedano, Nathan A Baertsch

**Affiliations:** Seattle Children’s Research Institute

## Abstract

Breathing is an essential motor behavior that emerges from the activity of rhythm-generating neurons in the preBötzinger Complex (preBötC). Although the properties of preBötC neurons are sufficient for rhythmogenesis, their activity is continually shaped by long-range inputs that convey homeostatic, emotional, volitional, and behavioral influences. Here, we provide a systematic survey of monosynaptic projections from excitatory (glutamatergic) and inhibitory (GABAergic) neurons across the mouse brain that target the preBötC. Using a dual recombinase-dependent retrograde tracing approach, we compile an atlas of afferent regions, highlighting the breadth of excitatory and inhibitory inputs that converge on this critical network. This atlas underscores the preBötC as a nexus where multimodal signals are integrated to regulate rhythm generation and patterning. Rather than a self-contained oscillator, the preBötC is a hub whose function is tuned by diverse long-range excitatory and inhibitory influences. By defining the anatomical substrates of this input, we lay a foundation for dissecting the functional roles of higher-order brain regions in shaping breathing across physiological, behavioral, and pathological contexts.

**SIGNIFICANCE STATEMENT:** Breathing is controlled by a small brainstem region called the preBötzinger Complex, which generates the rhythm for each breath. Many other brain areas influence this region, but the sources of these inputs—and whether they excite or inhibit breathing—have not been systematically mapped. Here, we present a detailed anatomical atlas identifying where excitatory and inhibitory signals to the preBötzinger Complex arise across the mouse brain. This resource provides a structural framework for understanding how breathing is shaped by sensory signals, behavior, and brain state.

## INTRODUCTION

The mechanisms that control breathing are surprisingly complex given the deceptive stereotypy of its function. This complexity reflects the deep integration of respiratory circuits with nearly every major function of the brain^1^. Like other vital processes, breathing operates automatically, governed by continuous reflexive mechanisms that maintain homeostasis through precise regulation of blood gases and pH^2^. But unlike other automatic functions, breathing is uniquely embedded within a wide range of non-homeostatic contexts. Breathing is effortlessly adapted to behaviors such as speech, swallowing, laughing, and crying^3^, flexibly accommodates volitional demands such as playing wind instruments or prolonged breath-holding, and is strongly modulated by state. Transitions between sleep and wakefulness, shifts in arousal, and exposure to stress, pain, or noxious stimuli all profoundly reshape breathing^4,5^. How the brain coordinates and integrates these diverse modes of control—homeostatic, behavioral, volitional, and emotional—remains poorly understood. Addressing this question requires not only an understanding of the functionally specialized respiratory rhythm- and pattern-generating circuits within the brain, but also the long-range interactions that embed these circuits within broader neural networks.

The circuits responsible for automatically generating and regulating the motor commands that drive breathing are confined to the brainstem—primarily localized to the ventrolateral medulla in a rostrocaudally oriented region known as the ventral respiratory column (VRC). The VRC contains heterogeneous rhythm- and pattern-generating microcircuits whose integration coordinates the sequential phases of respiratory motor output—inspiration, post-inspiration, and active expiration^6^. At its core lies the preBötzinger Complex (preBötC), the principal rhythmogenic site for inspiration^7^—the obligatory phase of mammalian breathing. This bilateral cluster of neurons in the ventrolateral medulla underlies the neural rhythmicity that ultimately drives activation of inspiratory muscles like the diaphragm and is therefore essential for the continuity of breathing from birth until death^8^.

While much work has focused on understanding the sufficiency of the preBötC for rhythm generation—unravelling the intrinsic properties and excitatory–inhibitory interactions of its constituent neurons^9–12^—this circuit does not operate in isolation. Consistent with breathing’s complex functional integration, the preBötC receives extensive monosynaptic projections from diverse brain regions^7^. Many of these arise from nearby brainstem nuclei specialized for respiratory reflexes, including chemosensory regions that detect changes in CO_2_/H⁺ or O_2_ mechanosensory pathways that convey vagal and pulmonary stretch feedback, and interoceptive circuits that monitor airway patency and respiratory effort^1,13^. In addition to these canonical homeostatic inputs, the preBötC also receives projections from regions not traditionally considered part of the respiratory control network, including limbic and hypothalamic structures involved in emotional processing, cortical areas implicated in volitional motor planning, and midbrain circuits linked to arousal and defensive behaviors^14^. This broad convergence highlights the anatomical substrates by which the preBötC integrates homeostatic feedback with emotional, behavioral, and volitional influences. The scope of these inputs has been examined previously using G-deleted rabies viral tracing to identify inputs to transcriptionally defined excitatory SST and inhibitory GlyT2 neurons in the preBӧtC. Notably, these distinct subpopulations receive largely parallel inputs—the same brain regions project to both excitatory and inhibitory preBötC neurons^14^—suggesting that functional specificity arises more from the nature of upstream signals rather than from differential targeting of downstream preBötC subtypes. While this approach provided important insights, limitations of rabies tracing^15^—such as relatively low efficiency and non-cell-type-specific labeling of afferents—leave questions unresolved, including the molecular identity and prevalence of the projecting populations.

Here, we assemble a comprehensive survey of the brain-wide excitatory and inhibitory projections to the preBötC. In the process, we demonstrate the utility of an efficient, dual-recombinase-dependent tracing strategy to specifically identify transcriptionally-defined populations that project to a target region—in this case the preBötC. We highlight regional origins, relative prevalence of excitatory versus inhibitory inputs, and how these projections align with known functions of the source regions. This projection atlas reinforces the preBötC not as an isolated rhythmogenic kernel but as an embedded hub within a broadly distributed network. By mapping the convergent excitatory and inhibitory influences that reach the preBötC, we provide an anatomical framework for understanding multimodal breathing regulation, linking survival-driven homeostatic control with volitional, behavioral, and affective modulation.

## RESULTS

To map input projections to the preBötC arising from glutamatergic or GABAergic neurons, we implemented a dual-recombinase, dual-transgenic strategy (Fig. 1A). “Driver” mice expressing Cre recombinase under the control of the *Vglut2* or *Vgat* promoter were used to target glutamatergic or GABAergic populations, respectively. These mice were crossed with Ai65 reporter mice carrying a *Rosa26*-tdTomato allele requiring both Cre and FlpO recombinases for activation^16^. In the resulting heterozygous offspring (*Vglut2*^Cre/+^; Ai65 or *Vgat*^Cre/+^; Ai65), tdTomato fluorescence was absent without FlpO expression (n=3), confirming the expected dual-recombinase dependence. Projection-specific labeling was achieved by unilateral injection of a retrograde AAV encoding FlpO recombinase (rgAAV-FlpO) into the preBötC. The virus is taken up by axon terminals and transported to somata of projecting neurons^17^, where FlpO recombination activates tdTomato expression in transcriptionally defined (Cre-expressing) inputs. To verify injection accuracy and distinguish local from long-range neurons, a Cre-dependent eYFP reporter (AAV-DIO-eYFP) was co-injected. Animals were excluded if eYFP expression was absent or spread beyond the preBötC. In successful injections (n=3 of 6 *Vglut2*; Ai65, and n=3 of 9 *Vgat*; Ai65), robust tdTomato expression revealed labeled neurons distributed across the hindbrain, midbrain, interbrain, and forebrain (Fig. 1B). The density and locations of these input neurons differed between excitatory and inhibitory populations, revealing distinct patterns of long-range connectivity with the preBötC.

**Figure 1.**
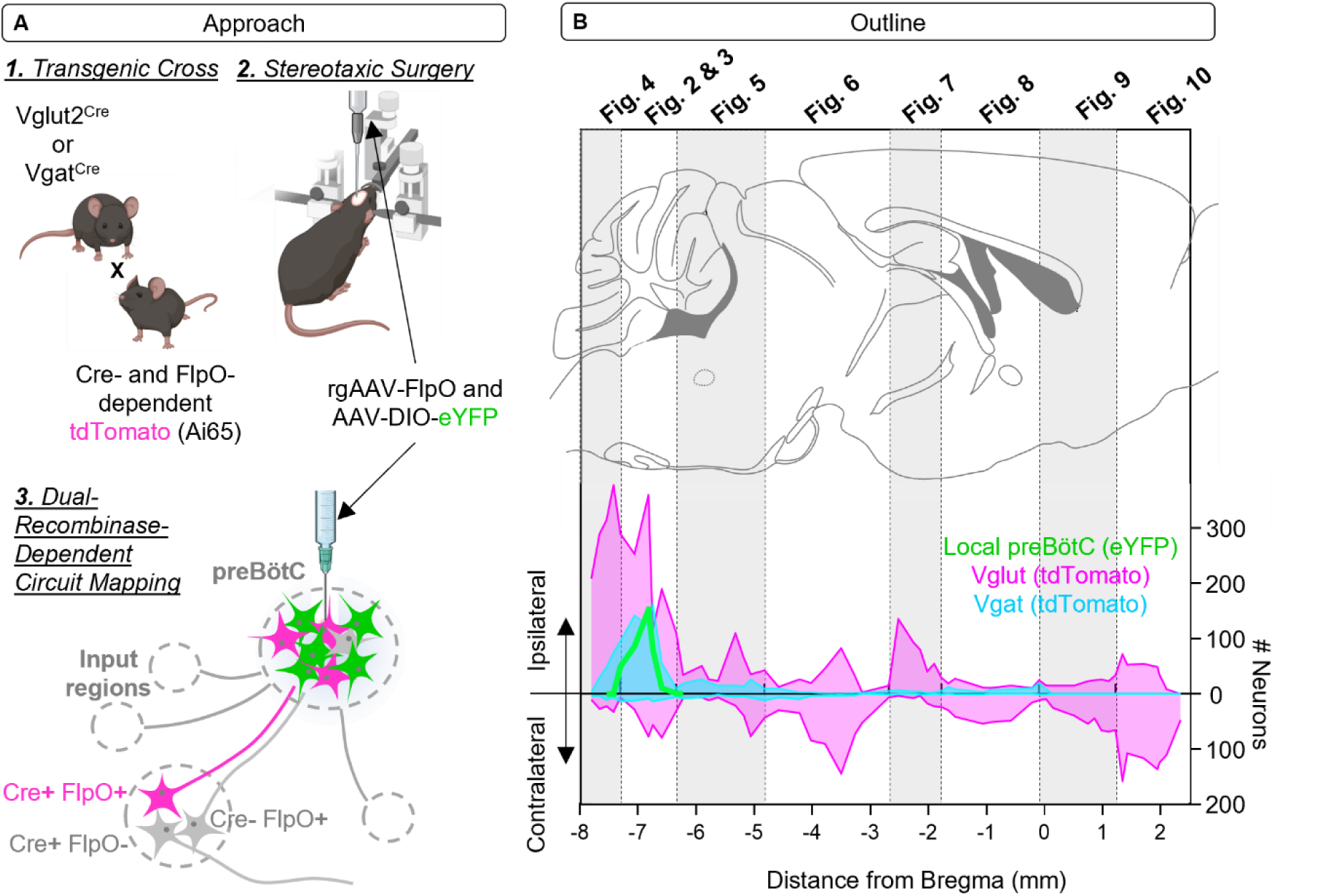
(**A**) Schematic of the dual-recombinase–dependent retrograde labeling strategy used to identify excitatory and inhibitory neurons projecting monosynaptically to the preBötC. An Ai65 (Cre- and FlpO-dependent tdTomato) reporter mouse was crossed with either Vglut2^Cre^ or Vgat^Cre^ driver lines. A mixed stereotaxic injection was performed in the preBötC consisting of a Cre-dependent GFP-expressing virus to mark the injection site (green) and a retrograde FlpO-expressing virus to label afferent neurons terminating in the preBötC. TdTomato expression (magenta) is restricted to neurons expressing both Cre and retrogradely delivered FlpO, thereby selectively labeling Vglut2^+^ or Vgat^+^ preBötC-projecting neurons. (**B**) Sagittal atlas reference sections from Paxinos and Franklin’s Mouse Brain Atlas illustrating the anteroposterior (Bregma) range represented in subsequent Figures. Plot shows counts of locally labeled preBötC neurons (green), Vglut2^+^ projection neurons (magenta), and Vgat^+^ projection neurons (cyan) as a function of Bregma position. Ipsilateral counts are plotted above the horizontal

### Local Connectivity

At the injection site, this approach delineated both local and commissural connectivity (Figs. 2 & 3). As expected, numerous glutamatergic and GABAergic input neurons were observed along the ipsilateral VRC^18,19^, extending rostrally and caudally from the injection site (Figs. 2A_1_-E_1_ & 3A_1_-E_1_), corresponding to the Bӧtzinger Complex and rostral ventral respiratory group (rVRG), respectively. Labeled glutamatergic neurons were also observed on the ventral medullary surface in the parafacial region, referred to as the retrotrepazoid nucleus or the parafacial respiratory group (RTN/pFRG; Fig. 2E_ii_)^20^. These local VRC inputs are core participants in the generation and dynamic shaping of rhythmic neuronal activity that supports robust and flexible breathing patterns^2,8,21^.

**Figure 2.**
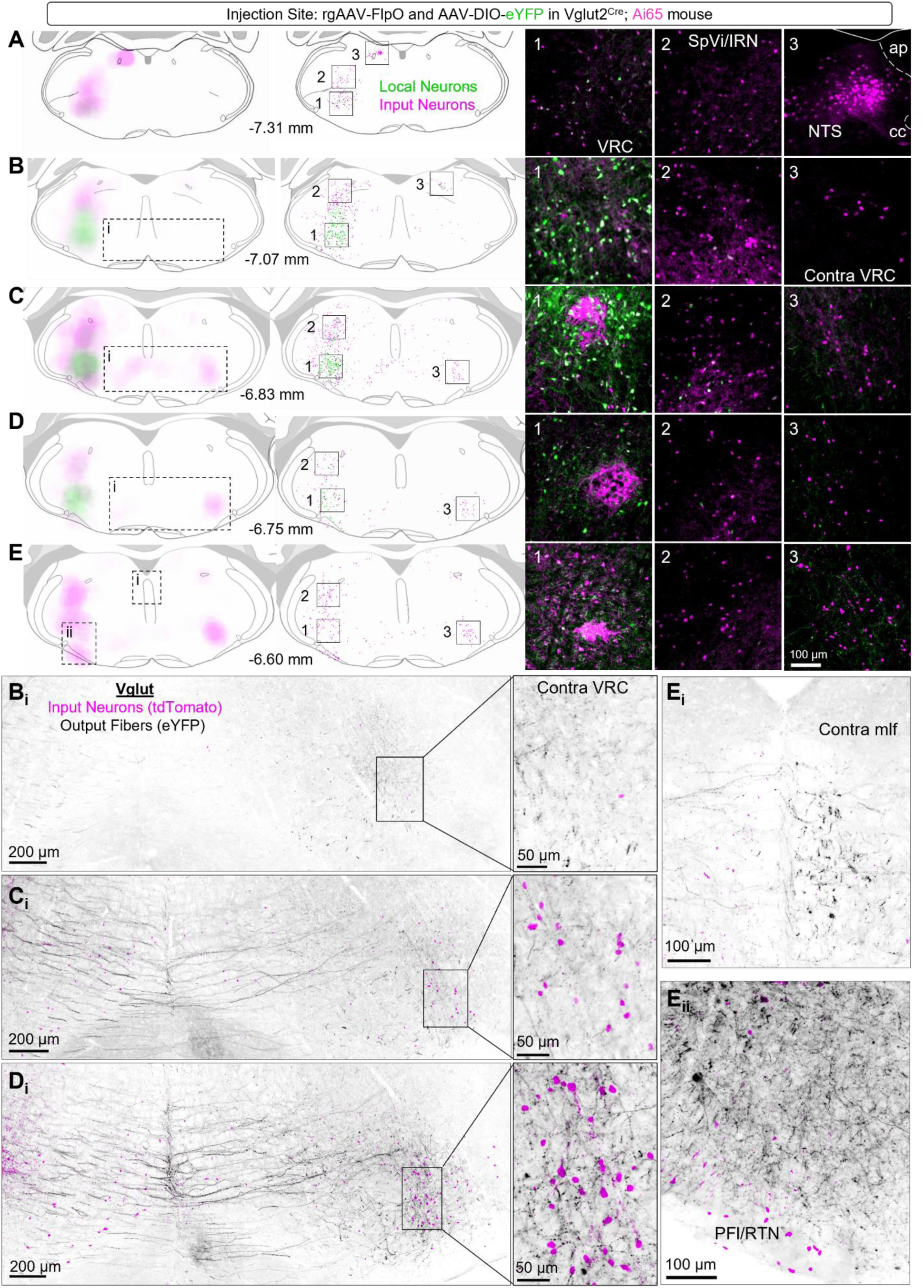
Peri-injection site labeling of excitatory projections to the preBötC in a Vglut2^Cre^; Ai65 mouse. **(A-E)** Pairs of coronal atlas reference sections shown at progressively rostral (ascending) Bregma levels. In each pair, the first column displays reference sections overlaid with density maps, and the second column displays the same sections overlaid with object plots indicating the locations of eYFP-labeled local neurons (green) and tdTomato-labeled excitatory input neurons (magenta). Boxed regions indicate areas shown at higher magnification in (1–3), highlighting local and preBötC-projecting neurons in pseudocolored fluorescence. Scale bar in E_3_ applies to all images. **(B_i_, C_i_, D_i_, E_i_, E_ii_)** Representative fluorescence images from a Vglut2^Cre^; Ai65 mouse showing output fibers labeled by eYFP fluorescence (grayscale) and monosynaptic input neurons labeled by tdTomato fluorescence (magenta). Image locations are indicated by dashed boxes on the corresponding reference sections. Callouts in B_i_, C_i_, and D_i_ highlight glutamatergic input neurons and output fibers in the contralateral ventral respiratory column (VRC). SpVi, interpolar part of the spinal trigeminal nucleus; IRN, intermediate reticular nucleus; NTS, nucleus tractus solitarius, PFl/RTN, lateral parafacial region/retrotrapezoid nucleus; mlf, medial longitudinal fascicle.

**Figure 3.**
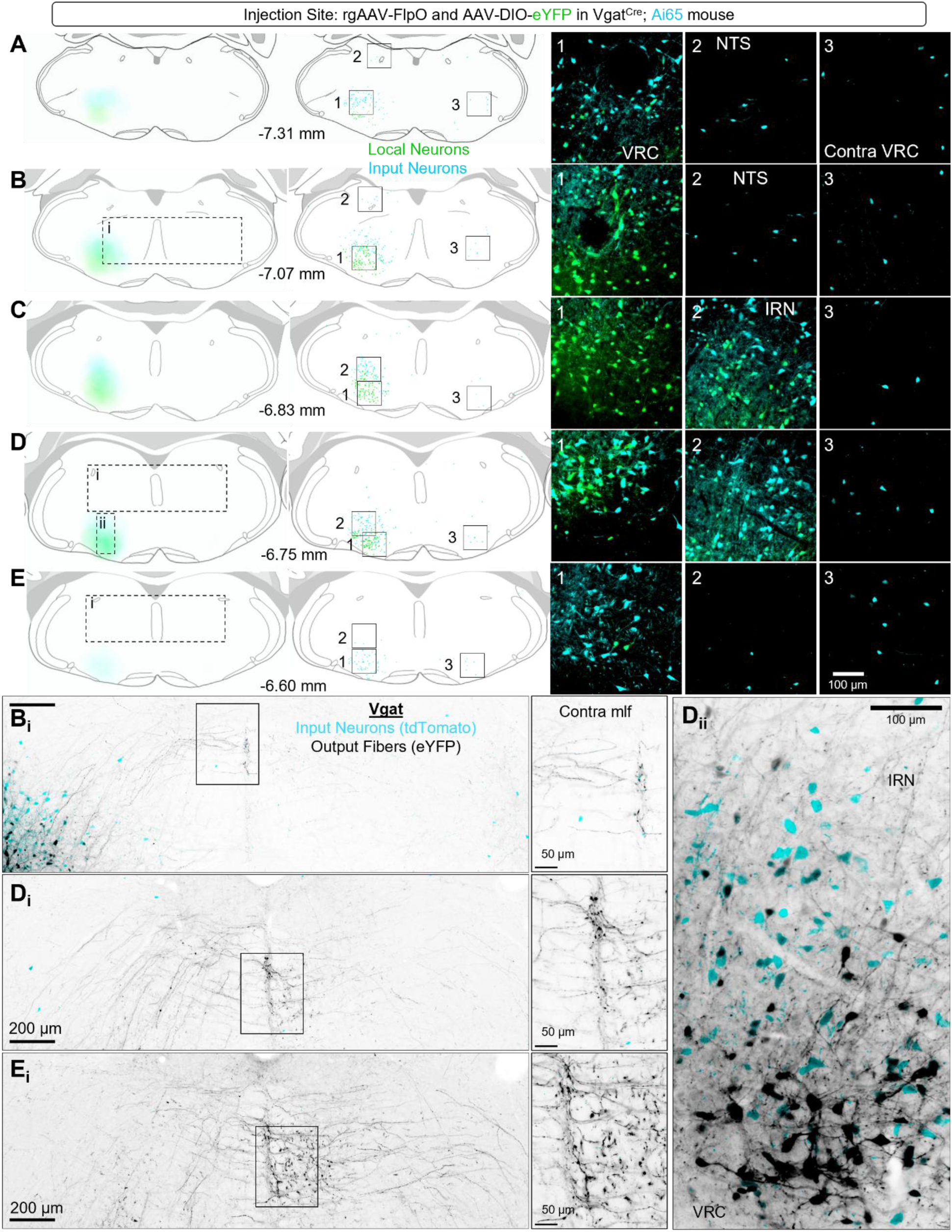
Peri-injection site labeling of inhibitory projections to the preBötC in a Vgat^Cre^; Ai65 mouse. **(A-E)** Pairs of coronal atlas reference sections shown at progressively rostral (ascending) Bregma levels. In each pair, the first column displays reference sections overlaid with density maps, and the second column displays the same sections overlaid with object plots indicating the locations of eYFP-labeled local neurons (green) and tdTomato-labeled inhibitory input neurons (cyan). Boxed regions indicate areas shown at higher magnification in (1–3), highlighting local and preBötC-projecting neurons in pseudocolored fluorescence. Scale bar in E_3_ applies to all images. **(B_i_, C_i_, D_i_, E_i_, E_ii_)** Representative fluorescence images from a Vgat^Cre^; Ai65 mouse showing output fibers labeled by eYFP fluorescence (grayscale) and monosynaptic input neurons labeled by tdTomato fluorescence (cyan). Image locations are indicated by dashed boxes on the corresponding reference sections. Boxes in B_i_, D_i_, and E_i_ correspond to magnified images to the right highlighting GABAergic output fibers in the medial longitudinal fascicle (mlf). VRC, ventral respiratory column; NTS, nucleus tractus solitarius; IRN, intermediate reticular nucleus

In addition to these local inputs, many input neurons were distributed outside the canonical VRC. Glutamatergic, but not GABAergic, neurons were scattered medial to the injection site (Fig. 2C, D), and extended dorsally into the intermediate reticular nucleus (IRN). A particularly notable density of glutamatergic input neurons was observed in the adjacent interpolar division of the spinal trigeminal nucleus (SpVi; Fig. 2A_2_-E_2_). Ipsilateral GABAergic input neurons were also found within the IRN but were largely restricted to regions immediately dorsal to the VRC and limited to the rostrocaudal extent of the injection site (Fig. 3B-D). Excitatory (Dbx1-expressing) neurons in the IRN dorsal to the preBötC have been identified as premotor neurons that drive inspiratory hypoglossal output^22^. The presence of reciprocal connectivity between these premotor regions and the preBötC supports bidirectional coupling between rhythmogenic and pattern-forming elements of the respiratory network, which may underlie interdependencies between respiratory rhythm and motor pattern generation^12^.

The presence of excitatory input neurons within the interpolar division of the SpV is notable given its role in processing orofacial somatosensory and proprioceptive signals and supporting sensorimotor integration^23,24^. SpVi neurons are engaged during active orofacial behaviors such as whisking and jaw movement and participate in coordinating facial sensory input with downstream motor patterning^25,26^. Our findings demonstrate direct synaptic input from SpVi neurons to the preBötC, indicating that trigeminal sensorimotor signals can access inspiratory circuitry without intermediary relays. Unlike VRC neurons, these inputs are unlikely to participate in rhythm generation but may provide a pathway through which ongoing orofacial behaviors dynamically modulate ventral respiratory column activity and adjust breathing pattern in a context-dependent manner.

Consistent with the established role of commissural excitation in synchronizing bilateral respiratory rhythm^27^, numerous glutamatergic input neurons were observed contralateral to the injection site, localized within the ventral IRN—providing a clear delineation of the VRC. These commissural glutamatergic neurons were positioned directly opposite the center of the injection site (Fig. 2C_3_); however, their highest density was skewed toward the rostral VRC (Fig. 2E_3_), with fewer labeled neurons in more caudal regions (Fig. 2A, B). This distribution suggests that excitatory commissural input to the preBötC arises preferentially from rostral VRC regions (Figs. 2A–E). In contrast, commissural GABAergic neurons were sparser and distributed more evenly along the VRC (Figs. 3A_3_-E_3_), consistent with weaker or more spatially diffuse inhibitory coupling across hemispheres^28^. The functional contribution of this inhibitory commissural coupling within the VRC remains to be determined.

eYFP labeling of local *Vgat*- and *Vglut2*-expressing preBötC neurons enabled visualization of their output projections. Although both populations targeted the contralateral VRC and dorsal motor regions including the hypoglossal motor pool (Figs. 2E_i_ & 3E_i_), their axons followed partially distinct trajectories. Commissural *Vglut2* axons predominantly coursed horizontally toward the midline, concentrated dorsal to the inferior olives, before entering rostrocaudally oriented fiber tracts within the ventral medial longitudinal fasciculus (mlf) (Fig. 2B_i_-D_i_). This trajectory mirrored the distribution of contralateral *Vglut2* input neurons, with ipsilateral fibers entering the mlf near the level of the injection site and exiting more rostrally to innervate the contralateral VRC. In contrast, *Vgat* axons preferentially followed a more dorsal route, targeting the midline mlf at the level of the hypoglossal (XII) motor nucleus before exiting to innervate the rostral VRC (Fig. 3B_i_, D_i_, E_i_), suggesting that excitatory and inhibitory preBötC neurons engage common medullary targets through anatomically distinct pathways.

Beyond local VRC targets, the output pathways of eYFP-labeled *Vglut2*- and *Vgat*-expressing preBötC neurons became increasingly distinct. Excitatory output fibers extended rostrally beyond the VRC into the parafacial region, where they were concentrated along the dorsal border of both the ipsilateral and contralateral facial (VII) motor nucleus (Fig. 5A_i_). These excitatory projections were largely non-overlapping with excitatory input pathways (tdTomato-labeled), suggesting that preBötC excitatory neurons engage downstream targets through projection patterns distinct from those providing input to the rhythmogenic network. In contrast, inhibitory output fibers were concentrated along the medial aspect of the ipsilateral VII nucleus (Fig. 5B_i_) and showed substantial overlap with inhibitory input pathways. This overlap raises the possibility that a subset of inhibitory fibers classified as “inputs” may arise from local preBötC neurons that collateralize to provide both local and long-range synaptic outputs. In addition to these projections, both excitatory and inhibitory output fibers were observed traversing dorsal regions of the contralateral gigantocellular reticular nucleus (GRN) (Fig. 6A_i_, B_i_). Because our labeling strategy was not optimized for anterograde visualization of synaptic terminals (e.g., via synaptophysin tagging), we cannot determine whether these fibers represent axons of passage or sites of synaptic contact. Moreover, fluorescence intensity diminished beyond these regions, precluding reliable tracing of output axons further rostrally.

### Other Medullary Inputs

Our tracing also revealed substantial input arising from medullary regions both caudal and rostral to the injection site. In the caudal medulla (Fig. 4), excitatory and inhibitory input neurons extended along the ventral medullary reticular nucleus (MDRNv) within a VRC region classically defined as the caudal ventral respiratory group (cVRG). Notably, inhibitory input neurons were prevalent ipsilateral to the injection site (Fig. 4B_i_), whereas excitatory input neurons were found predominantly in the contralateral cVRG (Fig. 4A-C). The cVRG is traditionally viewed as a premotor respiratory region enriched in expiratory-related neurons that project to spinal motor pools controlling abdominal and intercostal muscles^29^ and is recruited during conditions requiring active expiration such as hypercapnia, exercise, or airway challenge^30,31^. Input from this region to the preBötC therefore suggests reciprocal coupling between rhythm-generating and expiratory premotor compartments of the VRC, potentially enabling feedback from expiratory motor circuits to influence inspiratory timing or pattern under elevated respiratory demand.

**Figure 4.**
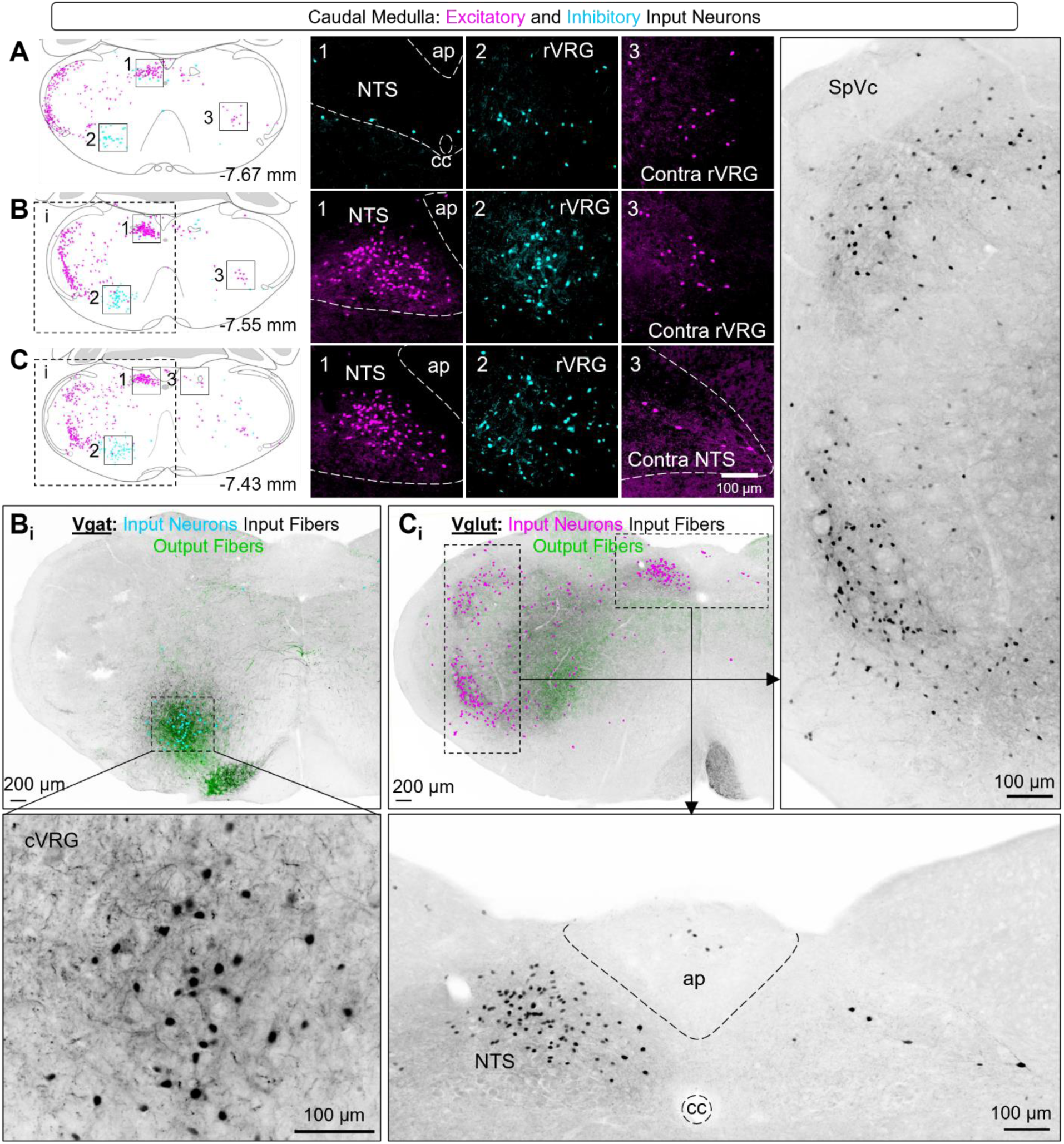
Excitatory and inhibitory inputs to the preBötC from the caudal medulla **(A-C)** Coronal reference sections shown at progressively rostral (ascending) Bregma levels with overlaid Vglut2⁺ (magenta) and Vgat⁺ (cyan) labeled neurons. Boxed regions indicate areas shown at higher magnification in (1–3), highlighting preBötC-projecting neurons in pseudocolored fluorescence. Scale bar in C_3_ applies to all images. **(B_i_)** Representative fluorescence image from a Vgat^Cre^; Ai65 mouse showing Vgat⁺ input fibers labeled by tdTomato fluorescence (grayscale), overlaid object plots indicating somata of Vgat⁺ neurons projecting to the preBötC (cyan), and local output fibers labeled by eYFP fluorescence (green). Image location is indicated by dashed boxes on the corresponding reference section. Callout shows grayscale tdTomato labeling in the caudal ventral respiratory group (cVRG). **(C_i_)** Representative fluorescence image from a Vglut2^Cre^; Ai65 mouse showing Vglut2⁺ input fibers labeled by tdTomato fluorescence (grayscale), overlaid object plots indicating somata of Vglut2⁺ neurons projecting to the preBötC (magenta), and local output fibers labeled by eYFP fluorescence (green). Image location is indicated by dashed boxes on the corresponding reference section. Callout shows grayscale tdTomato labeling in the spinal trigeminal nucleus caudalis (SpVc). rVRG, rostral ventral respiratory group; NTS, nucleus tractus solitarius; ap, area postrema; cc, central canal.

Extending beyond premotor respiratory regions, excitatory input neurons were also observed dorsally and ventrally within the caudal division of the spinal trigeminal nucleus (SpVc). The SpVc serves as the principal brainstem relay for nociceptive and thermal orofacial sensory information and is functionally analogous to the spinal dorsal horn^32^. In addition to direct projections to medullary respiratory circuits, SpVc neurons give rise to ascending projections to the parabrachial nucleus, providing an indirect route through which trigeminal nociceptive signals can engage autonomic and respiratory control networks^33^. Functionally, SpVc neurons are implicated in trigeminally driven defensive reflexes, including the mammalian diving response, in which facial sensory input produces rapid respiratory suppression and transient apnea^34,35^. Notably, this reflex persists following transection at the pontomedullary junction^36^, indicating that medullary circuits alone are sufficient to generate trigeminally evoked respiratory suppression. Consistent with this, noxious nasal or facial stimulation robustly engages SpVc while altering the activity of ventral medullary respiratory neurons^37^. Together, these findings indicate that excitatory SpVc neurons projecting directly to the preBötzinger Complex (preBötC) constitute a nociceptive trigeminal pathway capable of directly modulating inspiratory activity, enabling rapid suppression or reconfiguration of breathing during aversive or defensive states.

A dense population of excitatory input neurons was identified within the ipsilateral, and to a lesser extent the contralateral, nucleus of the solitary tract (NTS) (Fig. 2A-B & Fig. 4A-C). This observation is consistent with prior monosynaptic tracing studies in mice demonstrating that NTS neurons provide direct input to genetically defined preBötC populations^14^. In contrast, inhibitory NTS inputs were relatively sparse, comprising a small number of *Vgat*-expressing neurons distributed bilaterally within the NTS and often positioned along its ventral border relative to the bulk of excitatory NTS neurons (Fig. 3A_2_-B_2_ & Fig. 4A_1_). The NTS is a major integrative hub for visceral sensory afferents, including chemoreceptor and pulmonary stretch inputs, and plays a central role in shaping respiratory reflexes and phase transitions through its influence on ventral medullary respiratory circuits^38–40^. Together, the predominance of excitatory NTS input to the preBötC, coupled with a smaller inhibitory component, suggests that distinct NTS subpopulations may differentially modulate inspiratory rhythm and pattern in response to visceral sensory feedback, providing an anatomical substrate for reflexive adjustments of breathing.

Moving rostrally toward the pontomedullary junction, excitatory input neurons remained prominent within the spinal trigeminal nucleus, now concentrated in its oral division (SpVo) and the adjacent parvicellular reticular nucleus (PaRN). While both SpVo and SPVi process non-nociceptive trigeminal input, SpVo is more closely associated with premotor integration of orofacial and upper-airway behaviors, including jaw opening, mastication, and laryngeal control, rather than sensory reafference during active exploration^41^. Consistent with this role, the neighboring parvicellular reticular formation is a well-established premotor hub for jaw and laryngeal motor control and for coordinating patterned oropharyngeal movements^42,43^. The enrichment of excitatory preBötC input neurons in these rostral trigeminal–reticular regions therefore suggests a pathway specialized for coupling inspiratory timing to premotor orofacial and upper-airway motor programs, distinct from both defensive trigeminal pathways and the sensorimotor integration functions attributed to SPVi.

Inhibitory input neurons were also present within the magnocellular reticular nucleus (MARN; Fig. 2B–E), forming a triangular cluster surrounding the midline that was predominantly ipsilateral, with scattered contralateral neurons (Fig. 5B_1_, C_1_, E_1_). In contrast, excitatory input neurons in this region were sparse and largely restricted to more caudal portions of the MARN. This region, often referred to more broadly as the rostral ventral medulla (RVM), is well established as a key node for descending modulation of nociceptive processing and autonomic function^44–46^, integrating inputs from higher-order brain regions to coordinate behavioral, affective, and physiological responses to pain. Notably, manipulations of the RVM have also been shown to influence breathing; for example, local opioid signaling within the RVM contributes to opioid-induced respiratory depression^47^. The identification of inhibitory RVM neurons projecting directly to the preBötC therefore provides a potential anatomical substrate linking nociceptive and respiratory circuits, through which pain-related or analgesic signals may directly influence respiratory rhythm and pattern.

**Figure 5.**
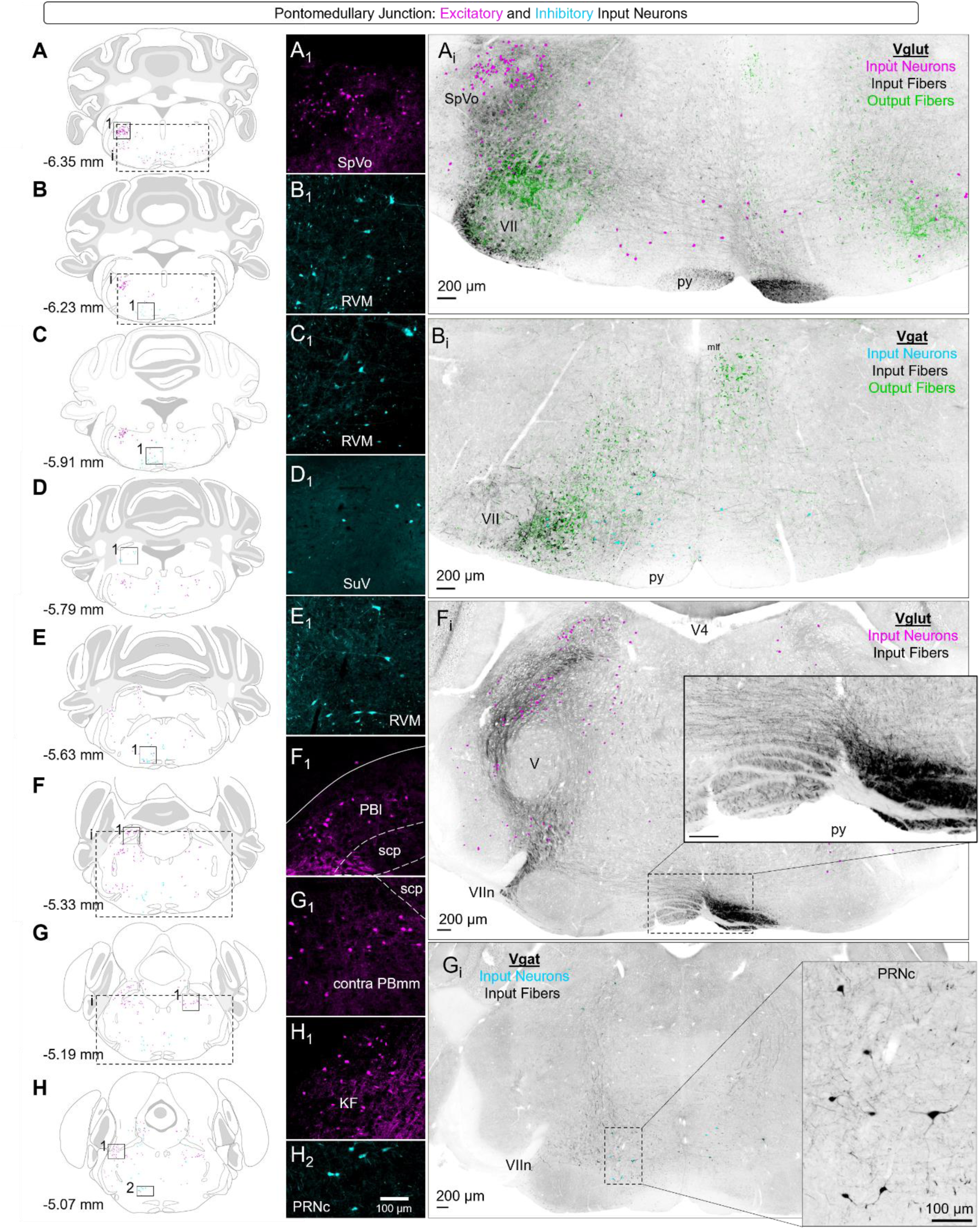
Excitatory and inhibitory inputs to the preBötC from the rostral medulla and pons **(A-H)** Coronal reference sections shown at progressively rostral (ascending) Bregma levels with overlaid Vglut2⁺ (magenta) and Vgat⁺ (cyan) labeled neurons. Boxed regions indicate areas shown at higher magnification in **(A_1_-H_1_, H_2_)** highlighting prominent clusters of preBötC-projecting neurons in pseudocolored fluorescence. Scale bar in H_2_ applies to all images. **(A_i_ & F_i_)** Representative fluorescence images from a Vglut2^Cre^; Ai65 mouse showing Vglut2⁺ input fibers labeled by tdTomato fluorescence (grayscale), overlaid object plots indicating somata of Vglut2⁺ neurons projecting to the preBötC (magenta), and local output fibers labeled by eYFP fluorescence (green). Image locations are indicated by dashed boxes on the corresponding reference sections. Callout in F_i_ shows grayscale tdTomato labeling of descending glutamatergic fibers in the pyramids (py) en route to the preBötC. **(B_i_ & G_i_)** Representative fluorescence image from a Vgat^Cre^; Ai65 mouse showing Vgat⁺ input fibers labeled by tdTomato fluorescence (grayscale), overlaid object plots indicating somata of Vgat⁺ neurons projecting to the preBötC (cyan), and local output fibers labeled by eYFP fluorescence (green). Image location is indicated by dashed boxes on the corresponding reference section. Callout in G_i_ shows grayscale tdTomato labeling of neurons in the pontine reticular nucleus caudalis (PRNc) that project to the preBötC. SpVo, oral part of spinal trigeminal nucleus; RVM, rostral ventral medulla; SuV, superior vestibular nucleus; PBl, lateral parabrachial nucleus; scp, superior cerebellar peduncle; PBmm, medial parabrachial nucleus; KF, Kölliker-Fuse; VII, facial motor nucleus; py, pyramids; V4, fourth ventricle; VIIn, facial nerve.

In the dorsal medulla, a smaller cluster of inhibitory input neurons was identified within the superior vestibular nucleus (SuV, Fig. 5D_1_). Although the vestibular nuclear complex is classically associated with postural and oculomotor control, it also engages autonomic and respiratory circuits through projections to the medullary reticular formation and ventral medulla^48^. Vestibular activation can modulate respiratory motor output and breathing pattern in rodents, particularly during movement or changes in body orientation^49,50^, and vestibular dysfunction in mice blunts ventilatory responses to hypercapnia^51^. The identification of inhibitory superior vestibular neurons projecting directly to the preBötC therefore suggests a potential substrate for vestibulo-respiratory coupling, through which postural or locomotor signals could suppress or reshape inspiratory activity.

In addition to labeling neuronal somata, our retrograde strategy yielded robust tdTomato expression in axonal fibers, enabling visualization of preBötC input pathways and collateral projections. Consistent with the predominance of excitatory inputs arising from midbrain and forebrain regions (see Fig. 1B), excitatory input fibers traversing the rostral medulla en route to the preBötC were abundant, whereas inhibitory input fibers were comparatively sparse and followed less distinct trajectories. Excitatory inputs were particularly dense within the contralateral—and to a lesser extent ipsilateral—pyramidal tract, with individual fibers observed exiting the pyramids and crossing the midline in the rostral medulla (Fig. 5A_i_, F_i_ inset) with collaterals continuing in the pyramidal tract through the caudal medulla (Fig. 4C_i_). Additional excitatory input fibers were prominent in the parafacial region, with the highest density along ventrolateral aspects of the medulla (Fig. 5A_i_), suggesting that some excitatory input neurons may collateralize to innervate both parafacial regions implicated in active expiration and the inspiratory preBötC. In contrast, inhibitory input fibers were absent from the pyramidal tract and instead preferentially coursed along the medial border of the facial (VII) motor nucleus, within a region corresponding to the lateral paragigantocellular reticular nucleus (PgRN) (Fig. 5B_i_).

### Pontine Inputs

The contribution of pontine structures to breathing has historically been difficult to resolve. Early transection and lesion studies showed that loss of pontine input profoundly alters respiratory patterning^52,53^, most notably causing apneusis, but these approaches lacked the spatial and cellular specificity required to assign distinct functions to defined pontine subregions across behavioral states. Recent advances, leveraging modern genetic and viral tools, have allowed for more nuanced dissection of pontine respiratory function, revealing complex, state- and context-dependent roles in shaping respiratory timing and pattern^3,54–58^. In this section, we describe both excitatory and inhibitory pontine substrates that are capable of directly modulating the preBötC.

Within the pontine central gray (PCG), a cluster of excitatory input neurons was observed in proximity to the locus coeruleus (LC) (Fig. 5E). The PCG is broadly implicated in processing sensory salience and affective valence, while the LC and peri-LC regions play central roles in regulating arousal, attention, and stress responses^59–62^. Excitatory projections from these regions to the preBötC therefore provide a plausible anatomical substrate through which changes in arousal or vigilance state could directly influence respiratory rhythm.

Rostral to the LC, additional excitatory input neurons were identified throughout the dorsal pons surrounding the superior cerebellar peduncle (scp), within regions collectively referred to as the parabrachial complex (PB) (Figs. 5F–H & 6A). The PB has long been implicated in the regulation of respiratory control but also serves as an integrative hub for many behaviors, affective states, and autonomic processes mediated by a rich diversity of transcriptionally-defined cell types^63,64^. Within the lateral PB (PBl), located dorsolateral to the scp, input neurons were predominantly ipsilateral, with the highest density found near the caudal PB (Fig. 5F_1_). In contrast, excitatory neurons in the medial PB (PBmm) were scattered bilaterally, with some neurons located in the adjacent caudal supratrigeminal nucleus (SuT; Fig. 5F_i_, G_1_). Moving rostrally, excitatory neurons became more clustered ventrolateral to the scp, particularly within the Kolliker Fuse nucleus (KF) (Fig. 5H_1_ & Fig. 6A_1_). The parabrachial complex as a whole is well known for its role in respiratory phase transitions, particularly in the termination of inspiration, and for its integration of sensory signals. More recent cell-type and projection specific manipulations implicate excitatory projections from the PBN/KF to the preBötC likely play a crucial role in providing excitatory drive to the preBӧtC, important for sustaining the homeostatic functions of breathing^65^, whereas glutamatergic subpopulations concentrated in the PBl, characterized by expression of tachykinin1 (Tac1) or calcitonin gene related peptide (CGRP), have opposing roles in regulating state-dependent non-homeostatic respiratory modulation during behaviors such as sniffing^54^. Thus, these direct excitatory inputs to the preBӧtC from different PB subdivisions likely reflect the multimodal nature of the parabrachial–preBötC circuit.

**Figure 6.**
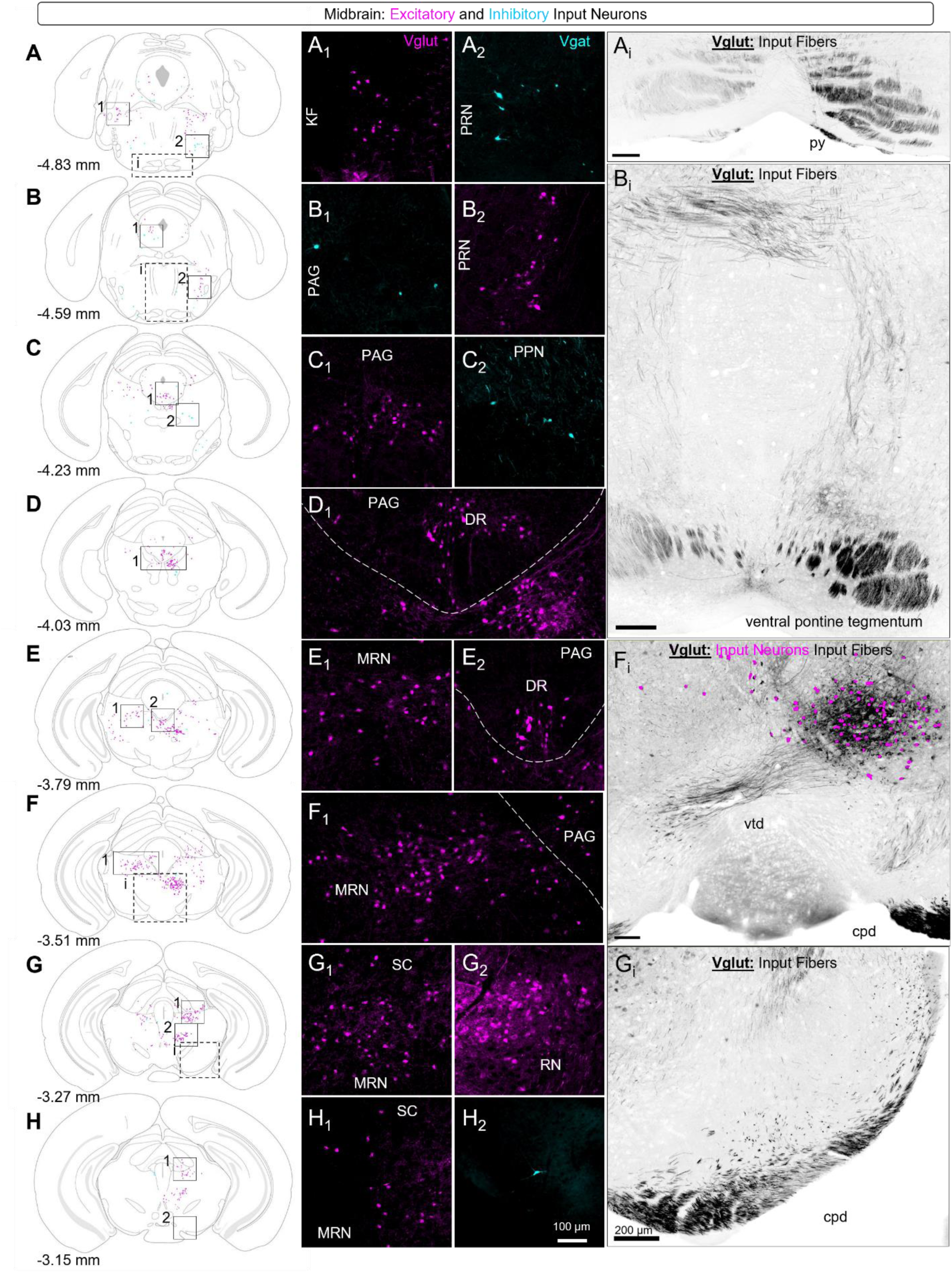
Excitatory and inhibitory inputs to the preBötC from the midbrain **(A-H)** Coronal reference sections shown at progressively rostral (ascending) Bregma levels with overlaid glutamatergic (magenta) and GABAergic (cyan) labeled neurons Boxed regions indicate areas shown at higher magnification in **(A_1_-H_2_)** highlighting prominent clusters of preBötC-projecting neurons in pseudocolored fluorescence. Scale bar in H_2_ applies to all inset images. **(A_i_, B_i_, F_i_, G_i_)** Representative fluorescence images from a Vglut2^Cre^; Ai65 mouse showing glutamatergic input fibers labeled by tdTomato fluorescence (grayscale) with overlaid object plots indicating somata of glutamatergic preBötC input neurons (magenta). Image locations are indicated by dashed boxes on the corresponding reference sections. KF, Kölliker-Fuse; PRN, pontine reticular nucleus; PAG, periaqueductal gray; DR, dorsal raphe; MRN, midbrain reticular nucleus; SC, superior colliculus; RN, red nucleus; py, pyramids; vtd, ventral tegmental decussation; cpd, cerebral peduncle.

In the ventral pons, scattered inhibitory input neurons were identified within the caudal pontine reticular nucleus (PRNc; Figs. 5F, G_i_, H_2_), consistent with prior reports demonstrating direct projections from this region to the preBötC^66^. These PRNc neurons have been implicated as a relay within corticobulbar pathways that support slow, regulated breathing and reduced negative affect, linking higher-order cortical activity to brainstem respiratory control^66^. This circuit has also been associated with behaviors requiring precise coordination of respiration with orofacial motor output, such as drinking. Direct inhibitory PRNc inputs to the preBötC therefore provide a potential substrate for top-down ponto-medullary gating of inspiratory activity, enabling suppression or slowing of breathing during coordinated orofacial behaviors and calm emotional states. More rostrally, inhibitory input neurons extended into lateral regions of the pontine reticular nucleus (Fig. 6A), where they became increasingly intermixed with a smaller population of excitatory input neurons (Fig. 6B_2_); the functional role of these mixed inputs remains unknown.

### Midbrain Inputs

At the level of the caudal midbrain, labeled input neurons were prominent in the periaqueductal gray (PAG) (Figs. 6B–F) and were predominantly excitatory, with a smaller population of inhibitory neurons distributed across PAG subdivisions. Caudally, excitatory input neurons were enriched within lateral and ventrolateral PAG regions (Fig. 6A, B), with labeling becoming progressively denser in ventral PAG domains at more rostral levels^67^ (Fig. 6C_1_–E_1_). At these rostral levels, clusters of excitatory input neurons were observed along the midline, corresponding anatomically to the dorsal raphe nucleus (DR; Fig. 6D_1_, E_1_). The PAG is widely recognized as a central integrative hub coordinating instinctive and emotional behaviors—including vocalization, defensive freezing, and behavioral apnea—via descending control of brainstem motor and autonomic circuits^68–71^. Consistent with prior monosynaptic tracing studies^14,67^, the direct PAG projections identified here provide a plausible anatomical substrate through which behavioral or emotional state signals could rapidly shape inspiratory activity.

Although the dorsal raphe nucleus is classically defined by its serotonergic neurons, many DR neurons are also capable of glutamate release, either alone or in combination with serotonin, enabling both fast excitatory and slower neuromodulatory influences on downstream targets^72,73^. Moreover, while the DR is often emphasized as part of ascending neuromodulatory systems, it also sends descending projections to medullary autonomic regions, providing a pathway through which midbrain serotonergic signaling can influence brainstem networks^74,75^. Through these descending projections, DR neurons are well-positioned to contribute to breathing stability, particularly across sleep–wake and arousal state transitions^76^.

Outside the periaqueductal gray, additional input neurons were observed within neighboring midbrain regions. Excitatory neurons, along with a small number of inhibitory neurons, were scattered throughout the ipsilateral midbrain reticular nucleus (MRN) lateral to the PAG (Fig. 6C, E_2_, F_1_), a region broadly associated with arousal and motor–autonomic integration. A few scattered excitatory input neurons extended dorsal of the MRN into the deep layer of the superior colliculus (SC) (Fig. 6F–H), a region implicated coordinating sensorimotor orienting behaviors^77^ and has been implicated in integrating visual and auditory cues with autonomic responses^78,79^—potentially linking orienting or startle behaviors with transient changes in breathing.

In more rostral regions of the midbrain, a dense cluster of excitatory input neurons was identified within the contralateral red nucleus (RN; Fig. 6F_i_, G_2_), consistent with prior monosynaptic tracing studies^14^. Although the RN is classically associated with fine motor coordination and limb movement control—and activation of RN neurons in mice evokes forelimb movements^80,81^—studies in neonatal and juvenile mammals indicate that the RN can also contribute to respiratory regulation under hypoxic conditions. In particular, lesions of the RN abolish the depressive phase of the biphasic hypoxic ventilatory response^82,83^, supporting a state-dependent pathway through which RN inputs may modulate inspiratory activity. Whether these preBötC-projecting RN neurons also participate in coordinating breathing with motor behaviors, such as locomotion, remains an open question.

Descending excitatory input fibers to the preBötC, or collaterals of these projecting neurons, were observed traversing the midbrain (Fig. 6A_i_, B_i_, F_i_, G_i_), whereas prominent inhibitory fiber tracts were minimal. Dense bundles of excitatory fibers were evident within the contralateral cerebral peduncle (cpd; Fig. 6G_i_) with fibers continuing through the ventral pontine tegmentum and basilar pons (Fig. 6A_i_, B_i_) before entering the medulla, consistent with corticoreticular and/or corticobulbar pathways in transit. A second prominent excitatory fiber tract was observed exiting the red nucleus, with axons crossing the midline via the ventral tegmental decussation (vtd; Fig. 6F_i_). Given that red nucleus outputs descend primarily via the rubrospinal tract, these labeled fibers may reflect collateralized motor-related pathways through which supramedullary excitatory inputs access preBötC circuitry.

### Interbrain Inputs

At the level of the diencephalon, labeled excitatory neurons wrapped around the medial edge of the cerebral peduncle, corresponding anatomically to the lateral hypothalamic area (LHA) and parasubthalamic nucleus (PSTN) (Fig. 7B-F). These excitatory input neurons were predominantly ipsilateral to the injection site, although a smaller contralateral population was observed at more rostral levels (Fig. 7E_1_). The LHA is a heterogeneous integrative hub involved in regulating arousal, motivation, and autonomic state, with orexin/hypocretin neurons playing a central role in narcolepsy and sleep-disordered breathing^84–86^. In contrast, the PSTN has been implicated in feeding suppression and integration of sensory/internal state signals^87,88^. Direct excitatory input from these excitatory interbrain neurons to the preBötC may support respiratory modulation during behavioral and internal state changes.

**Figure 7.**
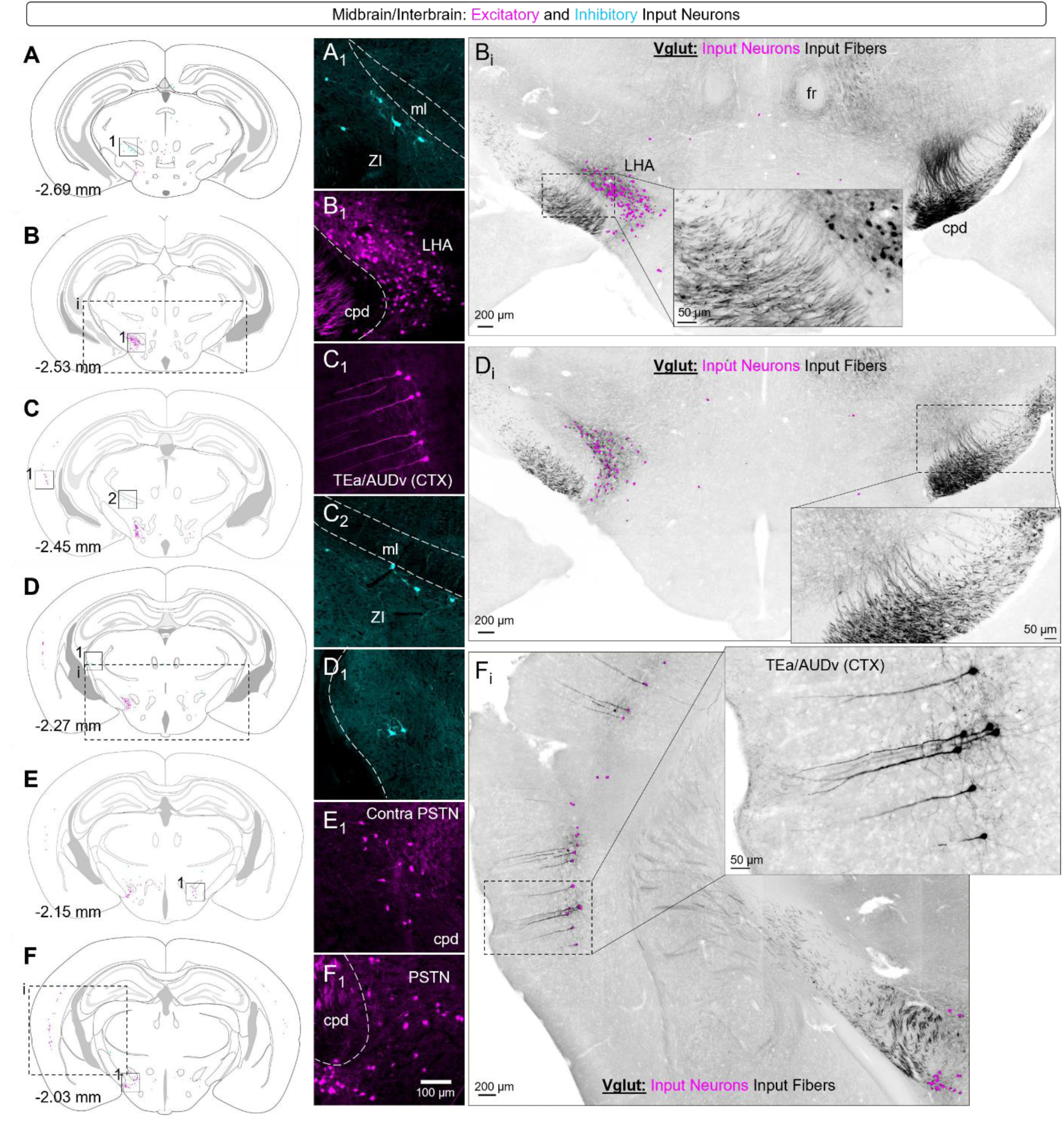
Excitatory and inhibitory inputs to the preBötC in the midbrain and interbrain **(A-F)** Coronal reference sections shown at progressively rostral (ascending) Bregma levels with overlaid glutamatergic (magenta) and GABAergic (cyan) labeled neurons. Boxed regions indicate areas shown at higher magnification in **(A_1_-F_1_)** highlighting prominent clusters of preBötC-projecting neurons in pseudocolored fluorescence. Scale bar in F_1_ applies to all inset images. **(B_i_, D_i_, F_i_)** Representative fluorescence images from a Vglut2^Cre^; Ai65 mouse showing glutamatergic input fibers labeled by tdTomato fluorescence (grayscale) with overlaid object plots indicating somata of glutamatergic preBötC input neurons (magenta). Image locations are indicated by dashed boxes on the corresponding reference sections. Callout in B_i_ shows grayscale tdTomato labeling of neurons in the lateral hypothalamic area (LHA) and descending glutamatergic fibers within the ipsilateral cerebral peduncle (cpd) that project toward the preBötC. Callout in D_i_ shows grayscale tdTomato labeling of fibers within the contralateral cerebral peduncle (cpd). Callout in F_i_ shows tdTomato labeling of gluatamatergic input neurons in temporal association/auditory (TEa/AUDv) cortex, highlighting prominent apical dendrites with complex branching into superficial cortical layers. ZI, zona incerta; ml, medial lemniscus; CTX, cortex; PSTN, parasubthalamic nucleus; fr, fasciculus retroflexus.

Dorsal to these regions, inhibitory input neurons were identified within the zona incerta (ZI), bordering the medial lemniscus (ml) (Fig. 7A–D). The ZI is a predominantly GABAergic structure with widespread descending projections to midbrain and brainstem motor centers, where it is thought to gate competing sensory and motor programs to support behavioral selection^89^. Direct inhibitory projections from the ZI to the preBötC therefore provide a plausible anatomical substrate through which inspiratory activity could be transiently suppressed, reshaped, or coordinated during motor behaviors that compete with breathing, including those engaging shared orofacial structures.

A discrete cluster of excitatory I”put ’eurons was identified within the ipsilateral dorsomedial hypothalamic nucleus (DMH), adjacent to the third ventricle near the midline (Fig. 8B_2_). The DMH is a key regulator of stress-evoked autonomic responses, including sympathetic activation and cardiovascular control, acting through descending projections to brainstem autonomic nuclei^90–92^. The presence of direct excitatory input from this region to the preBötC therefore suggests a pathway through which hypothalamic homeostatic signals may influence inspiratory activity. In contrast, no input neurons were detected within the arcuate hypothalamus, despite prior reports of arcuate projections to preBötC populations^14^, indicating that these previously described projection neurons may not be glutamatergic or GABAergic and thus were not captured by the present labeling strategy.

**Figure 8.**
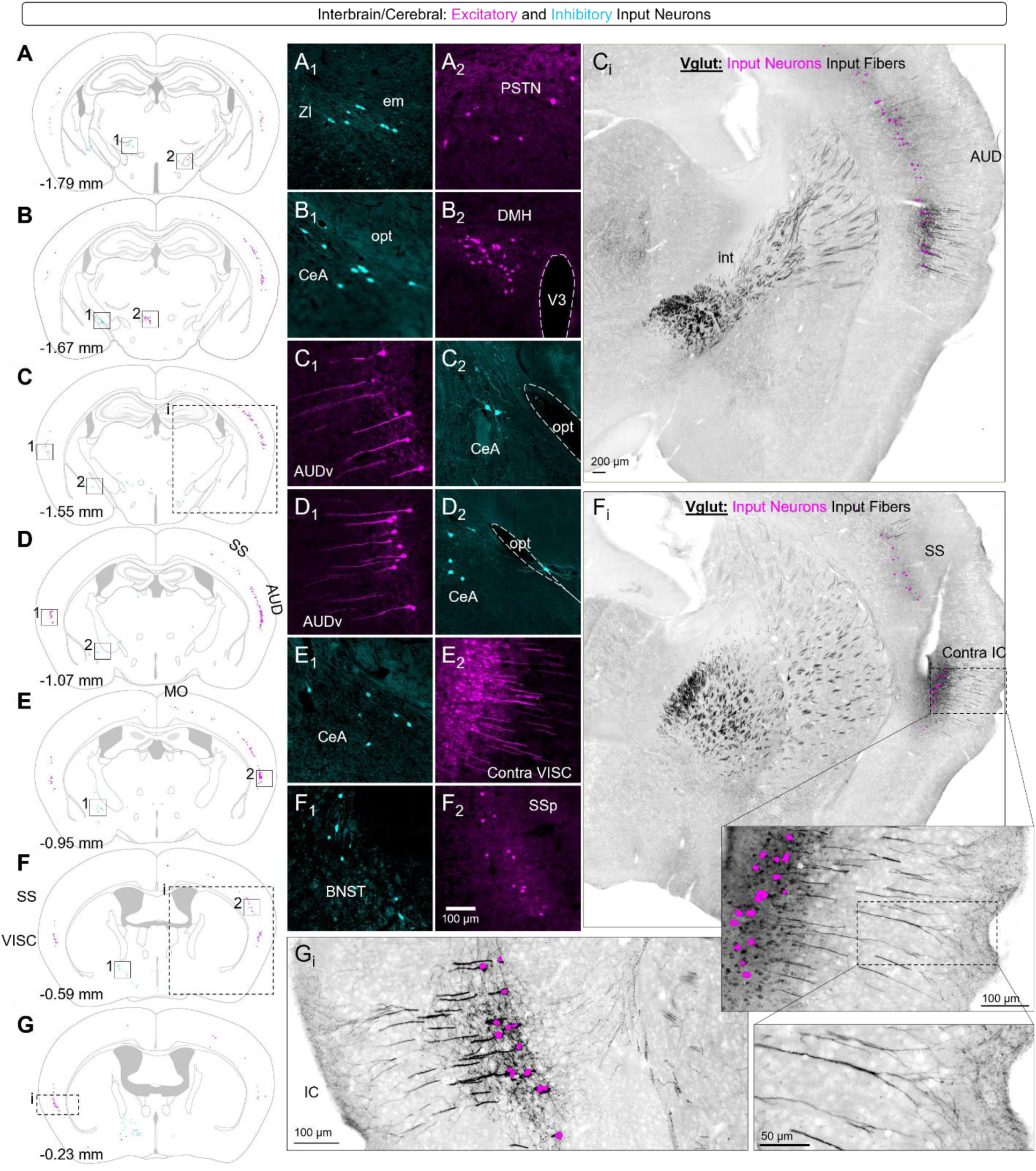
Excitatory and inhibitory inputs to the preBötC from the interbrain and cerebral cortex **(A-G)** Coronal reference sections shown at progressively rostral (ascending) Bregma levels with overlaid glutamatergic (magenta) and GABAergic (cyan) labeled neurons. Boxed regions indicate areas shown at higher magnification in **(A_1_-F_2_)** highlighting prominent clusters of preBötC-projecting neurons in pseudocolored fluorescence. Scale bar in F_2_ applies to all inset images. **(C_i_, F_i_, G_i_)** Representative fluorescence images from a Vglut2^Cre^; Ai65 mouse showing glutamatergic input fibers labeled by tdTomato fluorescence (grayscale) with overlaid object plots indicating somata of glutamatergic preBötC input neurons (magenta). Image locations are indicated by dashed boxes on the corresponding reference sections. Callout in F_i_ shows grayscale tdTomato labeling of neurons in the contralateral insular cortex (IC), highlighting prominent apical dendrites with complex branching into superficial cortical layers. ZI, zona incerta; em, external medullary lamina of the thalamus; PSTN, parasubthalamic nucleus; CeA, central amygdala; V3, third ventricle; DMH, dorsomedial nucleus of the hypothalamus; opt, optic tract; VISC, visceral area cortex; BNST, bed nucleus of the stria terminalis, SSp, primary somatosensory cortex; int, internal capsule.

### Cerebral Inputs

Within the cerebrum, the amygdala and bed nucleus of the stia terminalis (BNST), collectively referred to as the extended amygdala, regulates emotional reactivity, fear, and autonomic responses—triggering automatic survival responses to threat via their outputs to the hypothalamus, periaqueductal gray, and lower brainstem^93,94^. Inhibitory input neurons to the preBӧtC were found concentrated within the central subdivision of the amygdala (CeA) (Fig. 8B–E), consistent with prior reports and the largely GABAergic composition of this nucleus^95^. The CeA serves as a major output node of the amygdala, exerting downstream control over brainstem autonomic circuits. In humans, electrical stimulation of the amygdala can induce apnea^96^, supporting long-standing hypotheses that seizure spread to the amygdala may contribute to respiratory arrest in sudden unexpected death in epilepsy (SUDEP. However, optogenetic stimulations of CeA neurons in awake, behaving mice have revealed apparently conflicting effects, leading to respiratory acceleration rather than inducing apnea^97^. Farther rostral, a cluster of inhibitory input neurons, and a few scattered excitatory input neurons, were found in the ipsilateral bed nucleus of the stria terminalis (BNST) (Fig. 8F, G & 9A). Although direct modulation of respiratory rhythm by BNST neurons has not been definitively demonstrated, BNST involvement in CO_2_-evoked fear and anxiety^98^ suggests that BNST to preBӧtC circuits may alter respiratory control in response to threats such hypercapnia. Together, these projection pathways provide a plausible substrate through which extended amygdalar circuits may convey affective-, stress, or threat-related respiratory modulation.

Extensive labeling of excitatory neurons was observed bilaterally across several areas of the cortex (CTX). The majority of these cortical input neurons were located in layer 5 and exhibited large pyramidal somata with prominent apical dendrites extending toward superficial cortical layers (Fig. 7C_1_, F_i_), consistent with the morphology of corticofugal projection neurons. Layer 5 pyramidal neurons serve as the main output cells of the neocortex and give rise to long-range descending projections to subcortical and brainstem targets, including corticobulbar and corticoreticular pathways^99,100^. Consistent with prior descriptions of brainstem-projecting cortical neurons^101,102^, these cells were broadly represented across the cortical mantle, indicating that respiratory control may be accessed by multiple functional cortical domains rather than a single specialized region.

The most caudal clusters of cortical input neurons were located ipsilateral to the injection site within the temporal association area (TEa) and ventral auditory cortex (AUDv) of the lateral cortex (Fig. 7C_1_). Moving rostrally, labeled neurons expanded dorsally into primary and dorsal auditory cortical regions (AUDp/AUDd) (Fig. 7F_i_) and became increasingly prominent contralaterally, ultimately outnumbering ipsilateral auditory cortical neurons (Fig. 8A–C). These auditory and temporal association regions are involved in processing acoustic information relevant to communication, environmental salience, and orienting behaviors^103–105^. Direct inputs from these regions to the preBötC may provide a pathway for coordinating inspiratory timing with auditory-driven behavioral responses.

Moving rostrally, prominent excitatory input neurons emerged dorsally from auditory cortex into contralateral—though not ipsilateral—supplemental and primary somatosensory cortices (Fig. 8B–F), and subsequently into primary and secondary motor regions (MOp, MOs; Fig. 8E). At more rostral levels, excitatory input neurons within ipsilateral MOp/MOs became increasingly dense, with substantial contralateral populations also present (Figs. 9A–H and 10A–G). In many cases, labeled axons could be traced crossing the corpus callosum and external capsule and traversing the caudoputamen en route to the cerebral peduncle (Figs. 8C_i_, F_i_; 9C_i_, G_i_; 10A_i_), consistent with corticofugal projection pathways. Primary and secondary somatosensory cortices encode tactile and proprioceptive information related to body state and movement, while motor cortical regions are involved in the planning, initiation, and execution of voluntary actions, together forming an integrated sensorimotor network that supports coordinated behavior^106,107^. These distributed inputs from sensorimotor cortical regions may enable direct top-down modulation of inspiratory timing during voluntary breathing control as well as its automatic integration with other voluntary patterned motor behaviors such as locomotion or vocalization^57^.

**Figure 9.**
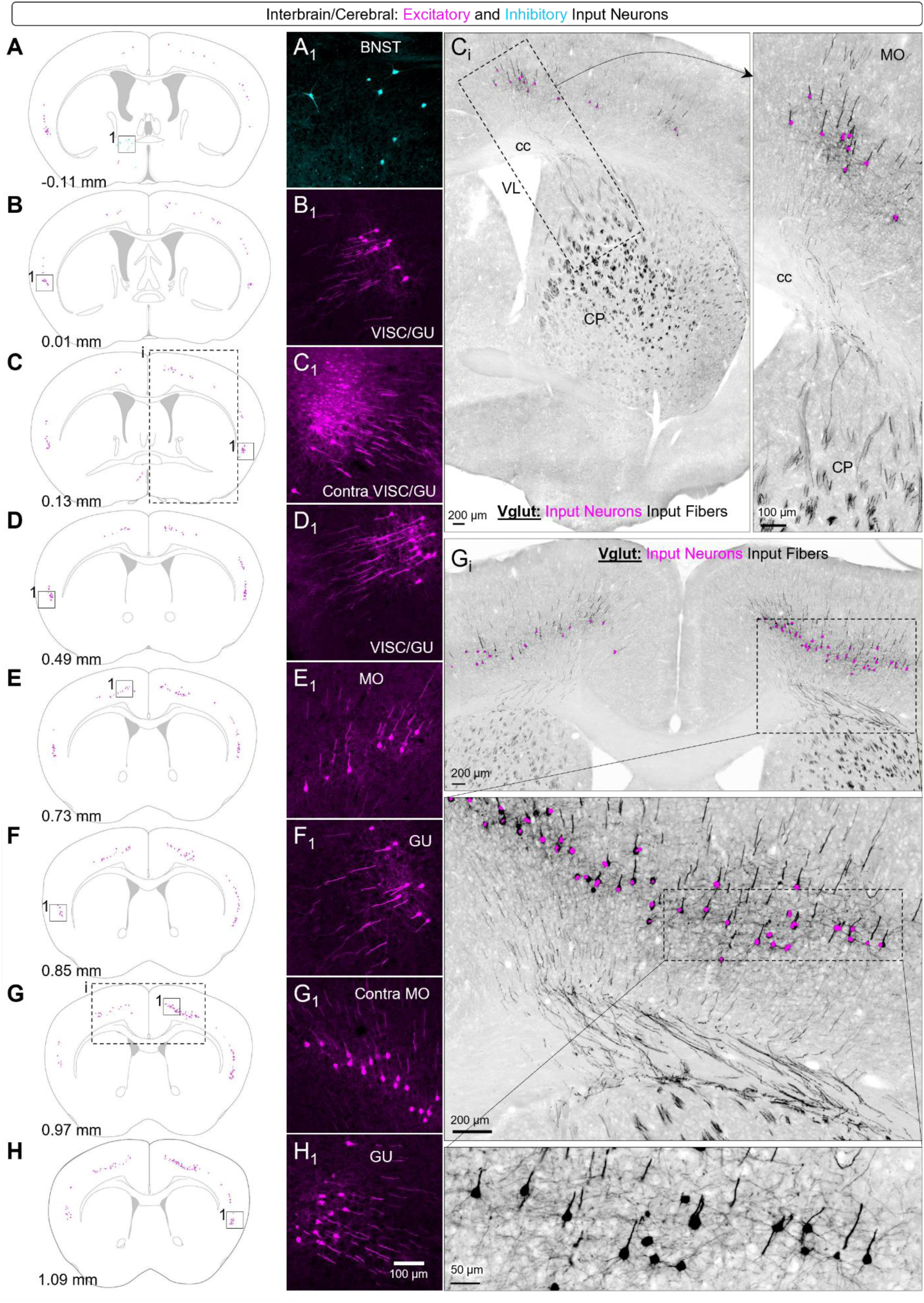
Excitatory and inhibitory inputs to the preBötC from the forebrain **(A-H)** Coronal reference sections shown at progressively rostral (ascending) Bregma levels with overlaid glutamatergic (magenta) and GABAergic (cyan) labeled neurons. Boxed regions indicate areas shown at higher magnification in **(A_1_-H_2_)** highlighting prominent clusters of preBötC-projecting neurons in pseudocolored fluorescence. Scale bar in H_1_ applies to all inset images. **(C_i_,G_i_)** Representative fluorescence images from a Vglut2^Cre^; Ai65 mouse showing glutamatergic input fibers labeled by tdTomato fluorescence (grayscale) with overlaid object plots indicating somata of glutamatergic preBötC input neurons (magenta). Image locations are indicated by dashed boxes on the corresponding reference sections. Callouts in C_i_ and G_i_ show grayscale tdTomato labeling of motor cortex (MO) neurons and their descending axons traversing the corpus collosum (cc) and entering the caudoputamen (CP) en route towards the preBötC. BNST, bed nucleus of the stria terminalis; VISC, visceral area of cortex; GU, gustatory area of cortex; VL, lateral ventricle.

In the lateral cortex, rostral to auditory regions, a well-defined bilateral cluster of excitatory input neurons to the preBötC was observed in the insula (Figs. 9A–H & 10A–C), concentrated near its visceral (VISC) and gustatory (GU) subdivisions (Figs. 9B_1_-D_1_, F_1_, H_1_ & 10A_1_). The insular cortex integrates interoceptive and emotional information, including awareness of breathing and dyspnea^108^. Human neuroimaging studies show that volitional breathing engages frontotemporal-insular regions^108^, while lesions to the right insula reduce perception of dyspnea and pain^109^. These findings suggest that insular projections may provide a direct pathway for adjusting preBötC excitability in response to internal state or perceived respiratory effort.

At the rostrocaudal level where the corpus callosum bifurcates, scattered excitatory neurons were observed within the infralimbic area (ILA) and anterior cingulate cortex (ACA) (Fig. 10Ci, D_1_, E). Electrical stimulation of the ILA in anesthetized rats suppresses expiratory-promoting reflexes while enhancing inspiratory-inhibitory responses, resulting in altered respiratory timing^110^. In the awake state, however, the ILA is a key node in emotional regulation and stress processing^111,112^, raising the possibility that its descending projections to the preBötzinger Complex (preBötC) provide a direct cortical pathway for top-down modulation of breathing during anxiety, affective state changes, or deliberate control of respiration. The ACA, by contrast, is more strongly associated with attentional control, interoceptive awareness, and affective processing^108,113^, suggesting that its projections may enable the integration of respiratory patterns with higher-order cognitive states such as conscious awareness or emotional appraisal.

**Figure 10.**
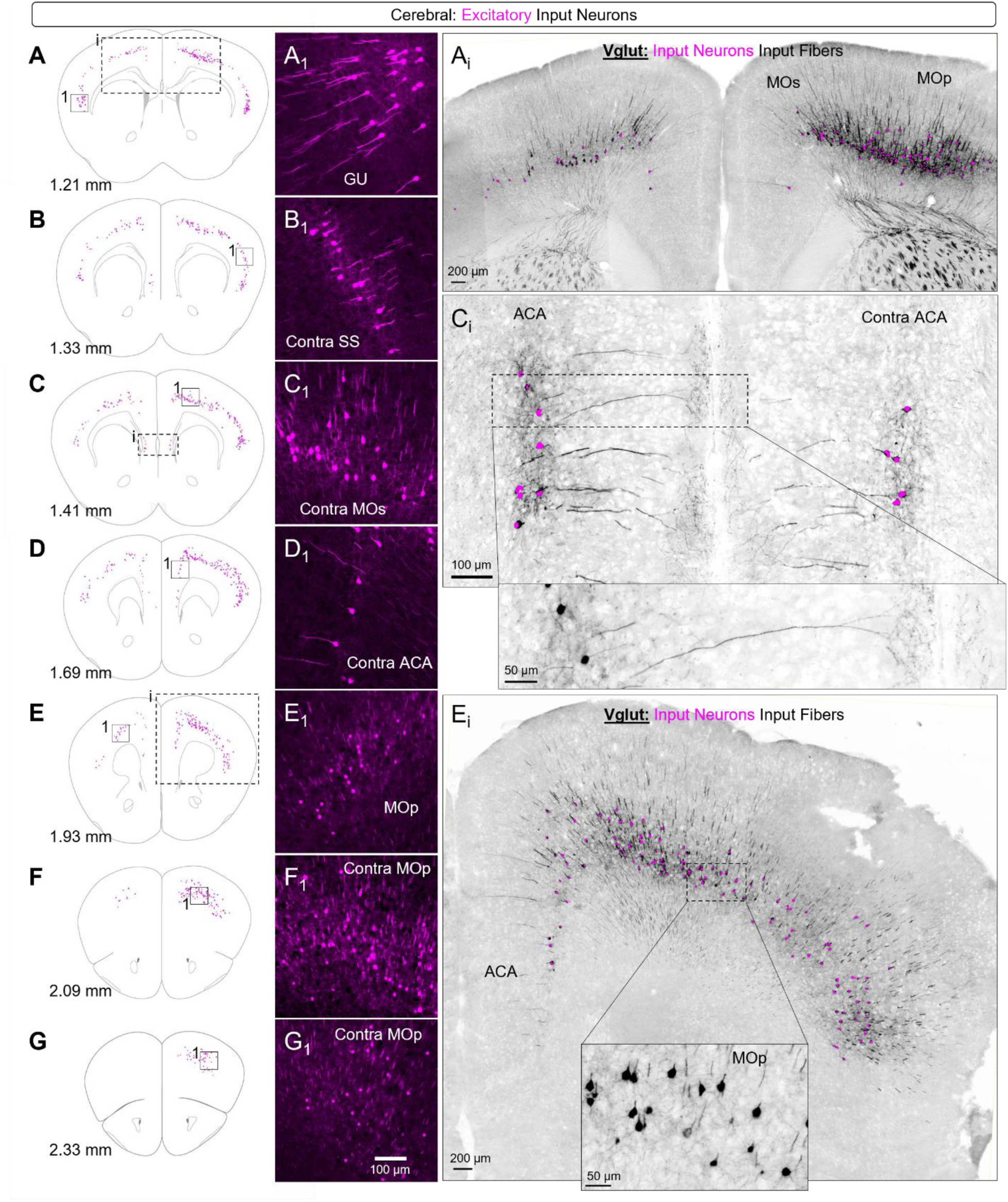
Excitatory and inhibitory inputs to the preBötC from the cerebral cortex. **(A-G)** Coronal reference sections shown at progressively rostral (ascending) Bregma levels with overlaid glutamatergic (magenta) labeled neurons. Boxed regions indicate areas shown at higher magnification in **(A_1_-G_1_)** highlighting prominent clusters of preBötC-projecting neurons in pseudocolored fluorescence. Scale bar in G_1_ applies to all inset images. **(A_i_, C_i_,E_i_)** Representative fluorescence images from a Vglut2^Cre^; Ai65 mouse showing glutamatergic input fibers labeled by tdTomato fluorescence (grayscale) with overlaid object plots indicating somata of glutamatergic preBötC input neurons (magenta). Image locations are indicated by dashed boxes on the corresponding reference sections. Callouts in C_i_ and E_i_ show grayscale tdTomato labeling of neuron morphology in anterior cingulate area (ACA) and primary motor (MOp) cortex regions, respectively. GU, gustatory area of cortex; SS, somatosensory cortex; MOs, secondary motor cortex.

At the rostrocaudal level where the corpus callosum bifurcates, scattered excitatory neurons were observed within the infralimbic area (ILA) (Fig. 10C_i_, D_1_, E) and anterior cingulate cortex (ACA) (Fig. 10D_1_, E). Electrical stimulation of the ILA in anesthetized rats suppresses expiratory-promoting reflexes while enhancing inspiratory-inhibitory responses, resulting in altered respiratory timing^110^. Beyond these reflexive effects, the ILA is a key node in emotional regulation and stress processing in the awake state^111^, raising the possibility that its descending projections to the preBötC provide a direct cortical pathway for top-down modulation of breathing during anxiety, affective state changes, or deliberate control of respiration. In contrast, the ACA is more strongly associated with attentional control, interoceptive awareness, and affective processing^108,113^. Its engagement during tasks requiring cognitive monitoring of internal state suggests that ACA projections to respiratory circuits may support the integration of breathing with higher-order cognitive processes, such as conscious awareness of respiration^114^ or emotion-linked modulation of respiratory patterns.

Together, these findings indicate that excitatory cortical inputs to the preBötC are widespread, bilateral, and functionally diverse. Notably, cortical influences from both regions are likely to be strongly state-dependent, exerting minimal impact during unconscious states but becoming prominent during wakefulness, emotional challenge, or cognitively engaged breathing. Although direct functional evidence for their specific respiratory roles remains limited, these anatomical pathways are well positioned to mediate volitional, sensory, and emotional influences that reshape preBötC-driven respiratory rhythms.

## DISCUSSION

The preBötC serves as the essential inspiratory rhythmogenic site within the ventral respiratory column^6,9^, yet its activity is profoundly shaped by a distributed network of inputs spanning the brainstem to the cerebrum^14^. Our dual-recombinase retrograde tracing strategy, leveraging *Vglut2*- or *Vgat*-Cre drivers with AAVrg-FlpO, provides a comprehensive atlas of these monosynaptic projections distinguished by their excitatory or inhibitory neurotransmitter phenotype. This approach reveals dense local connectivity, prominent homeostatic reflex pathways, and extensive supramedullary inputs from regions implicated in sensory integration, nociception, emotion, arousal, and volitional control. These findings reinforce the preBötC not as an isolated oscillator but as a convergent hub that dynamically integrates homeostatic demands with non-homeostatic behavioral and cognitive influences, aligning with emerging frameworks of multimodal respiratory control^3,57^. Rather than reiterating region-by-region anatomy, here we group the identified inputs into functional domains that map onto major modes of breathing control.

### Rhythm and pattern generation

Local and commissural inputs within the ventrolateral medulla underpin the core architecture of a bilaterally synchronized inspiratory oscillator embedded within a longitudinal pattern-forming network (VRC)^19^. Excitatory coupling within and across hemispheres drives rhythm generation and ensures bilateral synchronization of respiratory output^10,27,28^, whereas local inhibitory inputs facilitate phase transitions and regulate network gain under varying physiological demands^11,115^. Projections from adjacent reticular zones, including the intermediate reticular nucleus (IRN) and broader ventral respiratory group (VRG), enable reciprocal interactions between rhythmogenic and premotor compartments^12^, supporting coordinated adjustments in inspiratory timing and motor pattern output during multifaceted respiratory and orofacial behaviors.

### Homeostatic reflex control

Brainstem sensory–integrative inputs, centered on the NTS and related medullary circuits, constitute the core homeostatic pathways that tune preBötC activity in response to visceral afference, chemosensory drive, pulmonary stretch feedback, and signals related to respiratory effort^116,117^. These excitatory inputs, accompanied by a smaller inhibitory population, are consistent with reflex circuitry that can amplify or suppress inspiratory drive^38,118^. In parallel, chemosensitive neurons in parafacial regions—namely the retrotrapezoid nucleus^13,119^—provide excitatory input capable of adjusting inspiratory drive in response to changes in CO_2_/H⁺. These medullary reflex pathways are further shaped by inputs from the parabrachial complex, which acts as an integrative relay linking sensory signals—many originating in the NTS—to downstream modulation of preBötC activity^120,121^. Together, these convergent inputs provide the anatomical foundation for continuous, reflexive stabilization of eupnea to maintain homeostasis across behavioral and arousal states.

### Pain, nociception, and defensive respiratory suppression

Direct monosynaptic inputs to the preBötC from medullary regions processing trigeminal nociceptive signals, together with inputs from brainstem pain-modulatory circuits, provide a pathway for rapid, forebrain-independent suppression or reconfiguration of inspiratory rhythm in response to aversive stimuli^37,47^. This circuitry supports defensive respiratory behaviors, including transient inspiratory pauses, protective apneas, and disruptions of breathing evoked by noxious or threatening stimulation of the face or upper airways^34^. In addition to trigeminal relays, the parabrachial nucleus serves as a key integrative hub for nociceptive and aversive signaling, relaying pain-related information to both forebrain and brainstem targets and contributing to the affective and autonomic dimensions of pain^122,123^. Importantly, the PB also projects directly to respiratory centers, providing a route through which nociceptive and aversive signals may rapidly influence preBötC activity^14,124^. In parallel, inhibitory projections from theRVM—a major node for descending nociceptive facilitation and inhibition^125^—offer an additional substrate through which ongoing pain states or analgesic processes may directly modulate breathing.

### Arousal- and state-dependent mode switching

Breathing is shaped by global brain state: transitions between sleep states and wakefulness, changes in arousal or vigilance, and pharmacological states such as anesthesia or opioid-induced sedation profoundly alter respiratory rhythm, variability, and responsiveness to sensory input^126–129^. Pontine and midbrain inputs—including the parabrachial complex, peri-locus coeruleus and pontine central gray regions, periaqueductal gray, and raphe-associated populations—are well positioned to contribute to this control via their projections to the preBötC. Collectively, these regions regulate arousal, stress, and defensive state, and their convergent projections offer multiple parallel routes through which global state signals may regulate the activity and neuromodulatory milieu of the preBötC network—ensuring breathing remains appropriately matched to behavioral and physiological context.

### Emotion and threat-linked control

Inhibitory projections from extended amygdala structures provide a direct pathway through which affective and threat-processing circuits may influence inspiratory rhythm. These inputs offer a plausible substrate for fear- and anxiety-associated respiratory changes, including transient inspiratory suppression, irregular breathing, or pattern reconfiguration during threat detection, as well as longer-lasting alterations in respiratory dynamics associated with sustained negative affect or anxiety states^96,97^. Beyond the amygdala itself, emotional modulation of breathing emerges from a broader network that integrates threat appraisal, interoceptive awareness, and autonomic control.

The periaqueductal gray (PAG) likely occupies a central position within this network, coordinating defensive behaviors, autonomic responses, and respiratory adjustments during fear, stress, and aversive contexts. PAG activation is closely linked to both behavioral apnea and changes in breathing pattern during defensive states, and it is strongly implicated in the affective perception of dyspnea^69,130^. In parallel, the insular cortex integrates interoceptive signals related to respiratory effort, air hunger, and bodily state^108,109^, contributing to the conscious perception of dyspnea and emotional salience of breathing. Descending projections from insular and medial prefrontal regions, including infralimbic (ILA) and anterior cingulate cortex (ACA), provide additional routes for emotional appraisal, stress regulation, and cognitive monitoring to directly bias brainstem respiratory circuits^110,114^. Together, these convergent pathways suggest that emotion-linked respiratory modulation arises from coordinated interactions among limbic, midbrain, and cortical systems that can directly access the preBötC.

### Volitional and behavioral integration

Descending corticofugal inputs indicate that the preBötC can be accessed by multiple sensorimotor cortical domains rather than a specialized “breathing cortex.” In particular, projections from primary and secondary somatosensory and motor cortices, together with auditory and temporal association areas, provide parallel routes for top-down modulation of inspiratory activity. Somatosensory and motor cortical inputs are well positioned to support volitional control of breathing and to coordinate inspiratory timing with ongoing or planned movements, while auditory cortical inputs may enable coupling of respiration with sound-guided and communicative behaviors^104^. Together, these pathways suggest that breathing is embedded within broader sensorimotor control loops, allowing inspiratory timing to be adjusted automatically in concert with learned or complex behaviors such as vocalization, orienting, or locomotion, without requiring continuous explicit control.

Importantly, cortical influences on breathing are integrated with midbrain hubs, particularly the periaqueductal gray (PAG), which coordinates innate behavioral and emotional responses. PAG activation is closely linked to both behavioral apnea and innate vocalization, and its descending projections to brainstem respiratory circuits provide a mechanism for translating sensorimotor cortical signals into rapid, stereotyped respiratory adjustments^67,70,101,130^. Through convergence of corticofugal and PAG-mediated pathways, voluntary intent, sensory context, and innate behavioral programs may jointly shape preBötC activity to ensure appropriate respiratory patterning during communication, defensive responses, and other sensorimotor behaviors.

### Advantages and caveats of the approach

This dual-recombinase strategy enables high-yield, transcriptionally constrained identification of projecting populations. By restricting reporter expression to neurons defined both by neurotransmitter identity and by projection to the injected site, this method provides a robust and interpretable anatomical map of preBötC-directed inputs with distinct molecular characteristics. However, labeling reflects projections to the injected region rather than exclusively to preBötC neurons and may therefore capture some afferents targeting immediately adjacent structures—such as the nucleus ambiguus or nearby ventrolateral medullary compartments. Moreover, projection labeling does not resolve the number of postsynaptic targets or synaptic strength, and therefore the number of labeled input neurons should not be interpreted as a reliable measure of circuit influence.

Additional limitations arise from the genetic constraints of the approach. By design, the analysis is restricted to *Vglut2*- and *Vgat*-expressing populations and may under-sample neuromodulatory or peptidergic inputs that fall outside these classes. Because Cre-mediated recombination is permanent, neurons that transiently express Cre during development remain susceptible to tdTomato labelling if FlpO is later expressed in adulthood. This likely explains the unexpectedly extensive *Vglut2*-dependent labeling observed in forebrain regions, despite the predominance of *Vglut1* expression in adult cortical pyramidal neurons: many cortical populations express *Vglut2* transiently during development before switching to *Vglut1*^131,132^, enabling persistent reporter expression following early recombination. Importantly, despite these considerations, glutamatergic and GABAergic projecting populations were clearly distinguished, allowing reliable comparison of excitatory and inhibitory inputs to the preBötC.

### Summary

Together, these findings support a distributed control architecture in which brainstem reflex circuits stabilize eupnea, local ventral respiratory column networks sustain rhythm and pattern coupling, nociceptive and defensive pathways enable rapid suppression or reconfiguration, pontine and midbrain hubs implement state-dependent mode switching, and forebrain and limbic systems provide direct access for volitional and affective modulation. In this framework, the preBötzinger Complex emerges as a central nexus where convergent excitatory and inhibitory signals repurpose a core rhythmogenic network to support diverse respiratory modes—from automatic homeostatic breathing to emotionally driven sighs and deliberate breath control. By defining the anatomical substrates of this integration, the present projection atlas provides a foundation for mechanistic dissection of how distributed brain networks shape breathing across behavioral states and offers a roadmap for understanding respiratory dysfunction in neurological and affective disorders.

## METHODS

### Animals

All experimental procedures were approved by the Seattle Children’s Research Institute (SCRI) Institutional Animal Care and Use Committee (IACUC; Protocol #ACUC00663) and were conducted in accordance with the NIH *Guide for the Care and Use of Laboratory Animals*. Transgenic mouse lines were obtained from The Jackson Laboratory and have been previously validated and described in the literature^133–135^. C57BL/6J mice were group housed on a 12-h light/dark cycle with food, water, and environmental enrichment available *ad libitum* in a temperature- and humidity-controlled vivarium at SCRI. Homozygous Ai65 mice (JAX #021875), carrying a Rosa26-tdTomato reporter allele requiring both Cre and FlpO recombinases for activation, were crossed with homozygous Vglut2-ires-Cre (JAX #028863) or Vgat-ires-Cre (JAX #028862) mice.

### Stereotaxic Viral Injections

At 6 weeks of age, heterozygous offspring from both genetic crosses received a unilateral 23 nL stereotaxic injection of rgAAV-EF1a-FlpO (Addgene #55637; Lot #v137041) mixed with AAV1-cDIO-eGFP (AAV1-hSyn-eGFP). Mice were anesthetized with inhaled isoflurane (1–3%), administered local lidocaine (s.c.) and ketoprofen (5 mg/kg, i.p.) for analgesia, and secured in a stereotaxic frame. Scalp hair was removed, and the skin was disinfected with alternating betadine and ethanol washes. A ∼2 cm midline incision was made to expose the skull. Fast Green FCF dye was applied to enhance visualization of cranial sutures, allowing the skull to be leveled and Bregma to be identified and zeroed. A small craniotomy was drilled above the targeted preBötC coordinates (AP: −6.8 mm; ML: ±1.3 mm). For each injection site, the dorsoventral coordinate (DV: −4.7 mm) was zeroed at the dura. A Nanoject syringe loaded with the viral mixture was then lowered to the target depth, and 23 nL was injected over 5 minutes before slow withdrawal of the pipette. The scalp was sutured, isoflurane anesthesia was discontinued, and mice were allowed to recover.

### Histology

Approximately 3 weeks after surgery, mice were deeply anesthetized with isoflurane and transcardially perfused with ice-cold phosphate-buffered saline (PBS; 40 mL), followed by ice-cold 4% paraformaldehyde (PFA) in PBS (40 mL). Brains were dissected and post-fixed in 4% PFA at 4 °C for 24 h, then transferred to 30% sucrose in PBS for cryoprotection. Once fully cryoprotected, brains were embedded in OCT mounting medium and stored at −80 °C until sectioning. Whole brains were sectioned coronally at 30 µm thickness using a cryostat. All sections were mounted onto glass slides and coverslipped with DAPI Fluoromount. Slides were imaged at 10× magnification using a V120 Olympus slide-scanning microscope. Sections encompassing the preBötC were imaged in the FITC channel to verify the injection site, and all sections were imaged in the TRITC channel to visualize labeled projections, as well as in the DAPI channel to provide anatomical landmarks.

### Image Processing

Injection site images from each brain were first examined in OlyVIA to determine inclusion. A representative *Vglut2*^Cre^; Ai65 and a representative *Vgat*^Cre^; Ai65 mouse were selected and used across all figures to illustrate the distribution of labeled neurons. Sections at matched Bregma levels from the selected *Vglut2*^Cre^; Ai65 and *Vgat*^Cre^; Ai65 brains were chosen and imported into QuPath projects, where contrast levels were adjusted to enhance neuron visibility while minimizing background signal. In sections containing labeled excitatory neurons, the tdTomato signal was pseudocolored magenta, whereas in sections containing labeled inhibitory neurons it was pseudocolored cyan. Regions containing representative clusters of labeled neurons were exported for use as figure callouts.

To generate atlas overlays, atlas images corresponding to the Bregma levels of the selected sections were modified to retain only contour lines. In QuPath, automated cell detection was performed on the DAPI channel to identify nuclei in each section. Detected cells were then classified based on tdTomato fluorescence using a supervised classification algorithm trained on representative image regions and manually verified for accuracy. Classified object images were spatially aligned to their corresponding atlas sections using BigWarp in ImageJ, with raw fluorescence images used as references for anatomical alignment. The *Vglut2* and *Vgat* object images were subsequently resized to match atlas dimensions and merged to generate composite projection maps. Figures were assembled in Microsoft Powerpoint.

## REFERENCES

1 Del Negro, C. A., Funk, G. D. & Feldman, J. L. Breathing matters. Nat Rev Neurosci 19, 351–367 (2018). 10.1038/s41583-018-0003-6

2 Feldman, J. L., Mitchell, G. S. & Nattie, E. E. Breathing: rhythmicity, plasticity, chemosensitivity. Annu Rev Neurosci 26, 239–266 (2003). 10.1146/annurev.neuro.26.041002.131103

3 Yackle, K. & Do, J. The multifunctionality of the brainstem breathing control circuit. Curr Opin Neurobiol 90, 102974 (2025). 10.1016/j.conb.2025.102974

4 Homma, I. & Masaoka, Y. Breathing rhythms and emotions. Exp Physiol 93, 1011–1021 (2008). 10.1113/expphysiol.2008.042424

5 Jafari, H., Courtois, I., Van den Bergh, O., Vlaeyen, J. W. S. & Van Diest, I. Pain and respiration: a systematic review. Pain 158, 995–1006 (2017). 10.1097/j.pain.0000000000000865

6 Ramirez, J. M. & Baertsch, N. A. The Dynamic Basis of Respiratory Rhythm Generation: One Breath at a Time. Annu Rev Neurosci 41, 475–499 (2018). 10.1146/annurev-neuro-080317-061756

7 Smith, J. C., Ellenberger, H. H., Ballanyi, K., Richter, D. W. & Feldman, J. L. Pre-Bötzinger complex: a brainstem region that may generate respiratory rhythm in mammals. Science 254, 726–729 (1991). 10.1126/science.1683005

8 Feldman, J. L. & Kam, K. Facing the challenge of mammalian neural microcircuits: taking a few breaths may help. J Physiol 593, 3–23 (2015). 10.1113/jphysiol.2014.277632

9 Rekling, J. C. & Feldman, J. L. PreBötzinger complex and pacemaker neurons: hypothesized site and kernel for respiratory rhythm generation. Annu Rev Physiol 60, 385–405 (1998). 10.1146/annurev.physiol.60.1.385

10 Ashhad, S. & Feldman, J. L. Emergent Elements of Inspiratory Rhythmogenesis: Network Synchronization and Synchrony Propagation. Neuron 106, 482–497.e484 (2020). 10.1016/j.neuron.2020.02.005

11 Baertsch, N. A., Baertsch, H. C. & Ramirez, J. M. The interdependence of excitation and inhibition for the control of dynamic breathing rhythms. Nat Commun 9, 843 (2018). 10.1038/s41467-018-03223-x

12 Ramirez, J. M. & Baertsch, N. Defining the Rhythmogenic Elements of Mammalian Breathing. Physiology (Bethesda) 33, 302–316 (2018). 10.1152/physiol.00025.2018

13 Guyenet, P. G. & Bayliss, D. A. Central respiratory chemoreception. Handb Clin Neurol 188, 37–72 (2022). 10.1016/B978-0-323-91534-2.00007-2

14 Yang, C. F., Kim, E. J., Callaway, E. M. & Feldman, J. L. Monosynaptic Projections to Excitatory and Inhibitory preBötzinger Complex Neurons. Front Neuroanat 14, 58 (2020). 10.3389/fnana.2020.00058

15 Patino, M. et al. Postsynaptic cell type and synaptic distance do not determine efficiency of monosynaptic rabies virus spread measured at synaptic resolution. Elife 12 (2023). 10.7554/eLife.89297

16 Madisen, L. et al. Transgenic mice for intersectional targeting of neural sensors and effectors with high specificity and performance. Neuron 85, 942–958 (2015). 10.1016/j.neuron.2015.02.022

17 Challis, R. C. et al. Adeno-Associated Virus Toolkit to Target Diverse Brain Cells. Annu Rev Neurosci 45, 447–469 (2022). 10.1146/annurev-neuro-111020-100834

18 Baertsch, N. A., Severs, L. J., Anderson, T. M. & Ramirez, J. M. A spatially dynamic network underlies the generation of inspiratory behaviors. Proc Natl Acad Sci U S A 116, 7493–7502 (2019). 10.1073/pnas.1900523116

19 Bush, N. E. & Ramirez, J. M. Latent neural population dynamics underlying breathing, opioid-induced respiratory depression and gasping. Nat Neurosci 27, 259–271 (2024). 10.1038/s41593-023-01520-3

20 Janczewski, W. A. & Feldman, J. L. Distinct rhythm generators for inspiration and expiration in the juvenile rat. J Physiol 570, 407–420 (2006). 10.1113/jphysiol.2005.098848

21 Anderson, T. M. & Ramirez, J. M. Respiratory rhythm generation: triple oscillator hypothesis. F1000Res 6, 139 (2017). 10.12688/f1000research.10193.1

22 Revill, A. L. et al. Dbx1 precursor cells are a source of inspiratory XII premotoneurons. Elife 4 (2015). 10.7554/eLife.12301

23 Moore, J. D., Kleinfeld, D. & Wang, F. How the brainstem controls orofacial behaviors comprised of rhythmic actions. Trends Neurosci 37, 370–380 (2014). 10.1016/j.tins.2014.05.001

24 Bosman, L. W. et al. Anatomical pathways involved in generating and sensing rhythmic whisker movements. Front Integr Neurosci 5, 53 (2011). 10.3389/fnint.2011.00053

25 Takatoh, J. et al. New modules are added to vibrissal premotor circuitry with the emergence of exploratory whisking. Neuron 77, 346–360 (2013). 10.1016/j.neuron.2012.11.010

26 Kurata, K. Hierarchical Organization Within the Ventral Premotor Cortex of the Macaque Monkey. Neuroscience 382, 127–143 (2018). 10.1016/j.neuroscience.2018.04.033

27 Bouvier, J. et al. Hindbrain interneurons and axon guidance signaling critical for breathing. Nat Neurosci 13, 1066–1074 (2010). 10.1038/nn.2622

28 Koizumi, H. et al. Structural-functional properties of identified excitatory and inhibitory interneurons within pre-Botzinger complex respiratory microcircuits. J Neurosci 33, 2994–3009 (2013). 10.1523/jneurosci.4427-12.2013

29 Iizuka, M. Respiration-related control of abdominal motoneurons. Respir Physiol Neurobiol 179, 80–88 (2011). 10.1016/j.resp.2011.01.003

30 Shiba, K., Siniaia, M. S. & Miller, A. D. Role of ventral respiratory group bulbospinal expiratory neurons in vestibular-respiratory reflexes. J Neurophysiol 76, 2271–2279 (1996). 10.1152/jn.1996.76.4.2271

31 Ezure, K., Tanaka, I. & Saito, Y. Brainstem and spinal projections of augmenting expiratory neurons in the rat. Neurosci Res 45, 41–51 (2003). 10.1016/s0168-0102(02)00197-9

32 Dubner, R. & Bennett, G. J. Spinal and trigeminal mechanisms of nociception. Annu Rev Neurosci 6, 381–418 (1983). 10.1146/annurev.ne.06.030183.002121

33 Cechetto, D. F., Standaert, D. G. & Saper, C. B. Spinal and trigeminal dorsal horn projections to the parabrachial nucleus in the rat. J Comp Neurol 240, 153–160 (1985). 10.1002/cne.902400205

34 Panneton, W. M. The mammalian diving response: an enigmatic reflex to preserve life? Physiology (Bethesda) 28, 284–297 (2013). 10.1152/physiol.00020.2013

35 Panneton, W. M. & Gan, Q. The Mammalian Diving Response: Inroads to Its Neural Control. Front Neurosci 14, 524 (2020). 10.3389/fnins.2020.00524

36 Panneton, W. M., Gan, Q. & Sun, D. W. Persistence of the nasotrigeminal reflex after pontomedullary transection. Respir Physiol Neurobiol 180, 230–236 (2012). 10.1016/j.resp.2011.11.012

37 Dutschmann, M. & Paton, J. F. Influence of nasotrigeminal afferents on medullary respiratory neurones and upper airway patency in the rat. Pflugers Arch 444, 227–235 (2002). 10.1007/s00424-002-0797-x

38 Bush, N. E., Oliveira, L. M., Glovak, Z. T. & Ramirez, J. M. An inspiration-off attractor supports the robust and flexible control of breathing. bioRxiv (2025). 10.1101/2025.09.23.678177

39 Feldman, J. L. & Del Negro, C. A. Looking for inspiration: new perspectives on respiratory rhythm. Nat Rev Neurosci 7, 232–242 (2006). 10.1038/nrn1871

40 Andresen, M. C. & Kunze, D. L. Nucleus tractus solitarius--gateway to neural circulatory control. Annu Rev Physiol 56, 93–116 (1994). 10.1146/annurev.ph.56.030194.000521

41 Ebert, C., Bagdasarian, K., Haidarliu, S., Ahissar, E. & Wallach, A. Interactions of Whisking and Touch Signals in the Rat Brainstem. J Neurosci 41, 4826–4839 (2021). 10.1523/JNEUROSCI.1410-20.2021

42 Travers, J. B., Yoo, J. E., Chandran, R., Herman, K. & Travers, S. P. Neurotransmitter phenotypes of intermediate zone reticular formation projections to the motor trigeminal and hypoglossal nuclei in the rat. J Comp Neurol 488, 28–47 (2005). 10.1002/cne.20604

43 Takatoh, J. et al. Constructing an adult orofacial premotor atlas in Allen mouse CCF. Elife 10 (2021). 10.7554/eLife.67291

44 Basbaum, A. I. & Fields, H. L. Endogenous pain control mechanisms: review and hypothesis. Ann Neurol 4, 451–462 (1978). 10.1002/ana.410040511

45 Fields, H. L., Heinricher, M. M. & Mason, P. Neurotransmitters in nociceptive modulatory circuits. Annu Rev Neurosci 14, 219–245 (1991). 10.1146/annurev.ne.14.030191.001251

46 Heinricher, M. M., Tavares, I., Leith, J. L. & Lumb, B. M. Descending control of nociception: Specificity, recruitment and plasticity. Brain Res Rev 60, 214–225 (2009).10.1016/j.brainresrev.2008.12.009

47 Phillips, R. S. et al. Pain-facilitating medullary neurons contribute to opioid-induced respiratory depression. J Neurophysiol 108, 2393–2404 (2012). 10.1152/jn.00563.2012

48 Balaban, C. D. & Porter, J. D. Neuroanatomic substrates for vestibulo-autonomic interactions. J Vestib Res 8, 7–16 (1998).

49 Xu, F., Zhuang, J., Zhou, T. R., Gibson, T. & Frazier, D. T. Activation of different vestibular subnuclei evokes differential respiratory and pressor responses in the rat. J Physiol 544, 211–223 (2002). 10.1113/jphysiol.2002.022368

50 Yates, B. J., Billig, I., Cotter, L. A., Mori, R. L. & Card, J. P. Role of the vestibular system in regulating respiratory muscle activity during movement. Clin Exp Pharmacol Physiol 29, 112–117 (2002). 10.1046/j.1440-1681.2002.03612.x

51 Allen, T. et al. Inner ear insult suppresses the respiratory response to carbon dioxide. Neuroscience 175, 262–272 (2011). 10.1016/j.neuroscience.2010.11.034

52 Alheid, G. F., Milsom, W. K. & McCrimmon, D. R. Pontine influences on breathing: an overview. Respir Physiol Neurobiol 143, 105–114 (2004). 10.1016/j.resp.2004.06.016

53 Dutschmann, M., Morschel, M., Kron, M. & Herbert, H. Development of adaptive behaviour of the respiratory network: implications for the pontine Kolliker-Fuse nucleus. Respir Physiol Neurobiol 143, 155–165 (2004). 10.1016/j.resp.2004.04.015

54 Arthurs, J. W., Bowen, A. J., Palmiter, R. D. & Baertsch, N. A. Parabrachial tachykinin1-expressing neurons involved in state-dependent breathing control. Nat Commun 14, 963 (2023). 10.1038/s41467-023-36603-z

55 Liu, S. et al. Divergent brainstem opioidergic pathways that coordinate breathing with pain and emotions. Neuron 110, 857–873.e859 (2022). 10.1016/j.neuron.2021.11.029

56 Lin, M. et al. Multiple Neural Networks Originating from the Lateral Parabrachial Nucleus Modulate Cough-like Behavior and Coordinate Cough with Pain. Am J Respir Cell Mol Biol 72, 272–284 (2025). 10.1165/rcmb.2024-0084OC

57 Baertsch, N. A., Reily, E., Sedano, J., Phillips, R. S. & Arthurs, J. W. Multimodal Breathing Control: Pontomedullary Mechanisms and Current Perspectives. Bioessays, e70072 (2025). 10.1002/bies.70072

58 Angelakos, C. C. et al. A cluster of neuropeptide S neurons regulates breathing and arousal. Curr Biol 33, 5439–5455.e5437 (2023). 10.1016/j.cub.2023.11.018

59 Benarroch, E. E. Locus coeruleus. Cell Tissue Res 373, 221–232 (2018). 10.1007/s00441-017-2649-1

60 Chen, X. Valence processing in pons. Neuron 111, 1353–1354 (2023). 10.1016/j.neuron.2023.04.009

61 Luskin, A. T. et al. Heterogeneous pericoerulear neurons tune arousal and exploratory behaviours. Nature 643, 437–447 (2025). 10.1038/s41586-025-08952-w

62 Xiao, C. et al. Glutamatergic and GABAergic neurons in pontine central gray mediate opposing valence-specific behaviors through a global network. Neuron 111, 1486–1503 e1487 (2023). 10.1016/j.neuron.2023.02.012

63 Pauli, J. L. et al. Molecular and anatomical characterization of parabrachial neurons and their axonal projections. Elife 11 (2022). 10.7554/eLife.81868

64 Nardone, S. et al. A spatially-resolved transcriptional atlas of the murine dorsal pons at single-cell resolution. Nat Commun 15, 1966 (2024). 10.1038/s41467-024-45907-7

65 Bateman, J. T. & Levitt, E. S. Opioid suppression of an excitatory pontomedullary respiratory circuit by convergent mechanisms. Elife 12 (2023). 10.7554/eLife.81119

66 Jhang, J., Park, S., Liu, S., O’Keefe, D. D. & Han, S. A top-down slow breathing circuit that alleviates negative affect in mice. Nat Neurosci 27, 2455–2465 (2024). 10.1038/s41593-024-01799-w

67 Trevizan-Bau, P. et al. Reciprocal connectivity of the periaqueductal gray with the ponto-medullary respiratory network in rat. Brain Res 1757, 147255 (2021). 10.1016/j.brainres.2020.147255

68 Deng, H., Xiao, X. & Wang, Z. Periaqueductal Gray Neuronal Activities Underlie Different Aspects of Defensive Behaviors. J Neurosci 36, 7580–7588 (2016). 10.1523/JNEUROSCI.4425-15.2016

69 Faull, O. K., Subramanian, H. H., Ezra, M. & Pattinson, K. T. S. The midbrain periaqueductal gray as an integrative and interoceptive neural structure for breathing. Neurosci Biobehav Rev 98, 135–144 (2019). 10.1016/j.neubiorev.2018.12.020

70 Holstege, G. The periaqueductal gray controls brainstem emotional motor systems including respiration. Prog Brain Res 209, 379–405 (2014). 10.1016/B978-0-444-63274-6.00020-5

71 Subramanian, H. H. & Holstege, G. Stimulation of the midbrain periaqueductal gray modulates preinspiratory neurons in the ventrolateral medulla in the rat in vivo. J Comp Neurol 521, 3083–3098 (2013). 10.1002/cne.23334

72 Gullino, L. S. et al. Evidence for a Role of 5-HT-glutamate Co-releasing Neurons in Acute Stress Mechanisms. ACS Chem Neurosci 15, 1185–1196 (2024). 10.1021/acschemneuro.3c00758

73 Wang, H. L. et al. Dorsal Raphe Dual Serotonin-Glutamate Neurons Drive Reward by Establishing Excitatory Synapses on VTA Mesoaccumbens Dopamine Neurons. Cell Rep 26, 1128–1142 e1127 (2019). 10.1016/j.celrep.2019.01.014

74 Bago, M., Marson, L. & Dean, C. Serotonergic projections to the rostroventrolateral medulla from midbrain and raphe nuclei. Brain Res 945, 249–258 (2002). 10.1016/s0006-8993(02)02811-1

75 Underwood, M. D., Arango, V., Bakalian, M. J., Ruggiero, D. A. & Mann, J. J. Dorsal raphe nucleus serotonergic neurons innervate the rostral ventrolateral medulla in rat. Brain Res 824, 45–55 (1999). 10.1016/s0006-8993(99)01181-6

76 Jacobs, B. L. & Azmitia, E. C. Structure and function of the brain serotonin system. Physiol Rev 72, 165–229 (1992). 10.1152/physrev.1992.72.1.165

77 May, P. J. The mammalian superior colliculus: laminar structure and connections. Prog Brain Res 151, 321–378 (2006). 10.1016/S0079-6123(05)51011-2

78 Keay, K. A., Redgrave, P. & Dean, P. Cardiovascular and respiratory changes elicited by stimulation of rat superior colliculus. Brain Res Bull 20, 13–26 (1988). 10.1016/0361-9230(88)90004-4

79 Lynch, E. et al. Descending pathways from the superior colliculus mediating autonomic and respiratory effects associated with orienting behaviour. J Physiol 600, 5311–5332 (2022). 10.1113/JP283789

80 Brovelli, A. et al. Dynamic Reconfiguration of Visuomotor-Related Functional Connectivity Networks. J Neurosci 37, 839–853 (2017). 10.1523/JNEUROSCI.1672-16.2016

81 Yeomans, J. S. & Tehovnik, E. J. Turning responses evoked by stimulation of visuomotor pathways. Brain Res 472, 235–259 (1988). 10.1016/0165-0173(88)90008-2

82 Ackland, G. L., Noble, R. & Hanson, M. A. Red nucleus inhibits breathing during hypoxia in neonates. Respir Physiol 110, 251–260 (1997). 10.1016/s0034-5687(97)00090-x

83 Waites, B. A., Ackland, G. L., Noble, R. & Hanson, M. A. Red nucleus lesions abolish the biphasic respiratory response to isocapnic hypoxia in decerebrate young rabbits. J Physiol 495 (Pt 1), 217–225 (1996). 10.1113/jphysiol.1996.sp021586

84 Bonnavion, P., Mickelsen, L. E., Fujita, A., de Lecea, L. & Jackson, A. C. Hubs and spokes of the lateral hypothalamus: cell types, circuits and behaviour. J Physiol 594, 6443–6462 (2016). 10.1113/JP271946

85 Sakurai, T. The neural circuit of orexin (hypocretin): maintaining sleep and wakefulness. Nat Rev Neurosci 8, 171–181 (2007). 10.1038/nrn2092

86 Varga, A. G., Whitaker-Fornek, J. R., Maletz, S. N. & Levitt, E. S. Activation of orexin-2 receptors in the Kӧlliker-Fuse nucleus of anesthetized mice leads to transient slowing of respiratory rate. Front Physiol 13, 977569 (2022). 10.3389/fphys.2022.977569

87 Kim, J. H. et al. A discrete parasubthalamic nucleus subpopulation plays a critical role in appetite suppression. Elife 11 (2022). 10.7554/eLife.75470

88 Shah, T., Dunning, J. L. & Contet, C. At the heart of the interoception network: Influence of the parasubthalamic nucleus on autonomic functions and motivated behaviors. Neuropharmacology 204, 108906 (2022). 10.1016/j.neuropharm.2021.108906

89 Chometton, S., Barbier, M. & Risold, P. Y. The zona incerta system: Involvement in attention and movement. Handb Clin Neurol 180, 173–184 (2021). 10.1016/B978-0-12-820107-7.00011-2

90 DiMicco, J. A., Samuels, B. C., Zaretskaia, M. V. & Zaretsky, D. V. The dorsomedial hypothalamus and the response to stress: part renaissance, part revolution. Pharmacol Biochem Behav 71, 469–480 (2002). 10.1016/s0091-3057(01)00689-x

91 Henderson, L. A. & Macefield, V. G. The role of the dorsomedial and ventromedial hypothalamus in regulating behaviorally coupled and resting autonomic drive. Handb Clin Neurol 180, 187–200 (2021). 10.1016/B978-0-12-820107-7.00012-4

92 Lowry, C. A., Plant, A., Shanks, N., Ingram, C. D. & Lightman, S. L. Anatomical and functional evidence for a stress-responsive, monoamine-accumulating area in the dorsomedial hypothalamus of adult rat brain. Horm Behav 43, 254–262 (2003). 10.1016/s0018-506x(02)00009-0

93 Lamotte, G., Shouman, K. & Benarroch, E. E. Stress and central autonomic network. Auton Neurosci 235, 102870 (2021). 10.1016/j.autneu.2021.102870

94 Schottelkotte, K. M. & Crone, S. A. Forebrain control of breathing: Anatomy and potential functions. Front Neurol 13, 1041887 (2022). 10.3389/fneur.2022.1041887

95 Gu, J., Sugimura, Y. K., Kato, F. & Del Negro, C. A. Central amygdala-to-pre-Botzinger complex neurotransmission is direct and inhibitory. Eur J Neurosci 60, 6799–6811 (2024). 10.1111/ejn.16589

96 Rhone, A. E. et al. A human amygdala site that inhibits respiration and elicits apnea in pediatric epilepsy. JCI Insight 5 (2020). 10.1172/jci.insight.134852

97 Wang, X. et al. GABAergic neurons in central amygdala contribute to orchestrating anxiety-like behaviors and breathing patterns. Nat Commun 16, 3544 (2025). 10.1038/s41467-025-58791-6

98 Taugher, R. J. et al. The bed nucleus of the stria terminalis is critical for anxiety-related behavior evoked by CO2 and acidosis. J Neurosci 34, 10247–10255 (2014). 10.1523/JNEUROSCI.1680-14.2014

99 Prasad, J. A., Carroll, B. J. & Sherman, S. M. Layer 5 Corticofugal Projections from Diverse Cortical Areas: Variations on a Pattern of Thalamic and Extrathalamic Targets. J Neurosci 40, 5785–5796 (2020). 10.1523/JNEUROSCI.0529-20.2020

100 Usrey, W. M. & Sherman, S. M. Corticofugal circuits: Communication lines from the cortex to the rest of the brain. J Comp Neurol 527, 640–650 (2019). 10.1002/cne.24423

101 Trevizan-Baú, P., Stanić, D., Furuya, W. I., Dhingra, R. R. & Dutschmann, M. Neuroanatomical frameworks for volitional control of breathing and orofacial behaviors. Respir Physiol Neurobiol 323, 104227 (2024). 10.1016/j.resp.2024.104227

102 Trevizan-Bau, P. et al. Forebrain projection neurons target functionally diverse respiratory control areas in the midbrain, pons, and medulla oblongata. J Comp Neurol 529, 2243–2264 (2021). 10.1002/cne.25091

103 Hackett, T. A. Information flow in the auditory cortical network. Hear Res 271, 133–146 (2011). 10.1016/j.heares.2010.01.011

104 King, A. J. & Schnupp, J. W. The auditory cortex. Curr Biol 17, R236–239 (2007). 10.1016/j.cub.2007.01.046

105 Souffi, S., Nodal, F. R., Bajo, V. M. & Edeline, J. M. When and How Does the Auditory Cortex Influence Subcortical Auditory Structures? New Insights About the Roles of Descending Cortical Projections. Front Neurosci 15, 690223 (2021). 10.3389/fnins.2021.690223

106 Aronoff, R. et al. Long-range connectivity of mouse primary somatosensory barrel cortex. Eur J Neurosci 31, 2221–2233 (2010). 10.1111/j.1460-9568.2010.07264.x

107 Lemon, R. N. Descending pathways in motor control. Annu Rev Neurosci 31, 195–218 (2008). 10.1146/annurev.neuro.31.060407.125547

108 Herrero, J. L., Khuvis, S., Yeagle, E., Cerf, M. & Mehta, A. D. Breathing above the brain stem: volitional control and attentional modulation in humans. J Neurophysiol 119, 145–159 (2018). 10.1152/jn.00551.2017

109 Schon, D. et al. Reduced perception of dyspnea and pain after right insular cortex lesions. Am J Respir Crit Care Med 178, 1173–1179 (2008). 10.1164/rccm.200805-731OC

110 Aleksandrov, V. G., Mercuriev, V. A., Ivanova, T. G., Tarasievich, A. A. & Aleksandrova, N. P. Cortical control of Hering-Breuer reflexes in anesthetized rats. Eur J Med Res 14 Suppl 4, 1–5 (2009). 10.1186/2047-783x-14-s4-1

111 Alexandrov, V. G., Ivanova, T. G. & Alexandrova, N. P. Prefrontal control of respiration. J Physiol Pharmacol 58 Suppl 5, 17–23 (2007).

112 Terreberry, R. R. & Neafsey, E. J. The rat medial frontal cortex projects directly to autonomic regions of the brainstem. Brain Res Bull 19, 639–649 (1987). 10.1016/0361-9230(87)90050-5

113 Dywan, J., Mathewson, K. J., Choma, B. L., Rosenfeld, B. & Segalowitz, S. J. Autonomic and electrophysiological correlates of emotional intensity in older and younger adults. Psychophysiology 45, 389–397 (2008). 10.1111/j.1469-8986.2007.00637.x

114 Peiffer, C., Poline, J. B., Thivard, L., Aubier, M. & Samson, Y. Neural substrates for the perception of acutely induced dyspnea. Am J Respir Crit Care Med 163, 951–957 (2001). 10.1164/ajrccm.163.4.2005057

115 Sherman, D., Worrell, J. W., Cui, Y. & Feldman, J. L. Optogenetic perturbation of preBötzinger complex inhibitory neurons modulates respiratory pattern. Nat Neurosci 18, 408–414 (2015). 10.1038/nn.3938

116 Zoccal, D. B., Furuya, W. I., Bassi, M., Colombari, D. S. & Colombari, E. The nucleus of the solitary tract and the coordination of respiratory and sympathetic activities. Front Physiol 5, 238 (2014). 10.3389/fphys.2014.00238

117 Ran, C., Boettcher, J. C., Kaye, J. A., Gallori, C. E. & Liberles, S. D. A brainstem map for visceral sensations. Nature 609, 320–326 (2022). 10.1038/s41586-022-05139-5

118 Deng, T. et al. A Molecularly Defined Medullary Network for Control of Respiratory Homeostasis. Adv Sci (Weinh) 12, e2412822 (2025). 10.1002/advs.202412822

119 Guyenet, P. G. et al. The Retrotrapezoid Nucleus: Central Chemoreceptor and Regulator of Breathing Automaticity. Trends Neurosci 42, 807–824 (2019). 10.1016/j.tins.2019.09.002

120 Chiang, M. C. et al. Parabrachial Complex: A Hub for Pain and Aversion. J Neurosci 39, 8225–8230 (2019). 10.1523/jneurosci.1162-19.2019

121 Campos, C. A., Bowen, A. J., Roman, C. W. & Palmiter, R. D. Encoding of danger by parabrachial CGRP neurons. Nature 555, 617–622 (2018). 10.1038/nature25511

122 Chen, Q. et al. Optogenetic Evidence for a Direct Circuit Linking Nociceptive Transmission through the Parabrachial Complex with Pain-Modulating Neurons of the Rostral Ventromedial Medulla (RVM). eNeuro 4 (2017). 10.1523/ENEURO.0202-17.2017

123 Roeder, Z. et al. Parabrachial complex links pain transmission to descending pain modulation. Pain 157, 2697–2708 (2016). 10.1097/j.pain.0000000000000688

124 Saper, C. B. & Loewy, A. D. Efferent connections of the parabrachial nucleus in the rat. Brain Res 197, 291–317 (1980). 10.1016/0006-8993(80)91117-8

125 De Preter, C. C. & Heinricher, M. M. The ’in’s and out’s’ of descending pain modulation from the rostral ventromedial medulla. Trends Neurosci 47, 447–460 (2024). 10.1016/j.tins.2024.04.006

126 Hao, X. et al. The Modulation by Anesthetics and Analgesics of Respiratory Rhythm in the Nervous System. Curr Neuropharmacol 22, 217–240 (2024). 10.2174/1570159X21666230810110901

127 Horner, R. L., Hughes, S. W. & Malhotra, A. State-dependent and reflex drives to the upper airway: basic physiology with clinical implications. J Appl Physiol (1985) 116, 325–336 (2014). 10.1152/japplphysiol.00531.2013

128 Vibert, J. F., Foutz, A. S., Caille, D. & Hugelin, A. Respiratory rhythm multistability during sleep-wake states. Brain Res 448, 403–405 (1988). 10.1016/0006-8993(88)91286-3

129 Zutler, M. & Holty, J. E. Opioids, sleep, and sleep-disordered breathing. Curr Pharm Des 17, 1443–1449 (2011). 10.2174/138161211796197070

130 Subramanian, H. H., Balnave, R. J. & Holstege, G. The midbrain periaqueductal gray control of respiration. J Neurosci 28, 12274–12283 (2008). 10.1523/JNEUROSCI.4168-08.2008

131 Boulland, J. L. et al. Expression of the vesicular glutamate transporters during development indicates the widespread corelease of multiple neurotransmitters. J Comp Neurol 480, 264–280 (2004). 10.1002/cne.20354

132 Wojcik, S. M. et al. An essential role for vesicular glutamate transporter 1 (VGLUT1) in postnatal development and control of quantal size. Proc Natl Acad Sci U S A 101, 7158–7163 (2004). 10.1073/pnas.0401764101

133 Daigle, T. L. et al. A Suite of Transgenic Driver and Reporter Mouse Lines with Enhanced Brain-Cell-Type Targeting and Functionality. Cell 174, 465–480 e422 (2018). 10.1016/j.cell.2018.06.035

134 Kouwenhoven, W. M. et al. VGluT2 Expression in Dopamine Neurons Contributes to Postlesional Striatal Reinnervation. J Neurosci 40, 8262–8275 (2020). 10.1523/JNEUROSCI.0823-20.2020

135 Vong, L. et al. Leptin action on GABAergic neurons prevents obesity and reduces inhibitory tone to POMC neurons. Neuron 71, 142–154 (2011). 10.1016/j.neuron.2011.05.028

